# A chemo-mechanical model of endoderm movements driving elongation of the amniote hindgut

**DOI:** 10.1101/2023.05.18.541363

**Authors:** Panagiotis Oikonomou, Helena C. Cirne, Nandan L. Nerurkar

## Abstract

While mechanical and biochemical descriptions of development are each essential, integration of upstream morphogenic cues with downstream tissue mechanics remains understudied in many contexts during vertebrate morphogenesis. A posterior gradient of Fibroblast Growth Factor (FGF) ligands generates a contractile force gradient in the definitive endoderm, driving collective cell movements to form the hindgut. Here, we developed a two-dimensional chemo-mechanical model to investigate how mechanical properties of the endoderm and transport properties of FGF coordinately regulate this process. We began by formulating a 2-D reaction-diffusion-advection model that describes the formation of an FGF protein gradient due to posterior displacement of cells transcribing unstable *Fgf8* mRNA during axis elongation, coupled with translation, diffusion, and degradation of FGF protein. This was used together with experimental measurements of FGF activity in the chick endoderm to inform a continuum model of definitive endoderm as an active viscous fluid that generates contractile stresses in proportion to FGF concentration. The model replicated key aspects of hindgut morphogenesis, confirms that heterogeneous - but isotropic - contraction is sufficient to generate large anisotropic cell movements, and provides new insight into how chemomechanical coupling across the mesoderm and endoderm coordinates hindgut elongation with outgrowth of the tailbud.

**Graphical Abstract:** 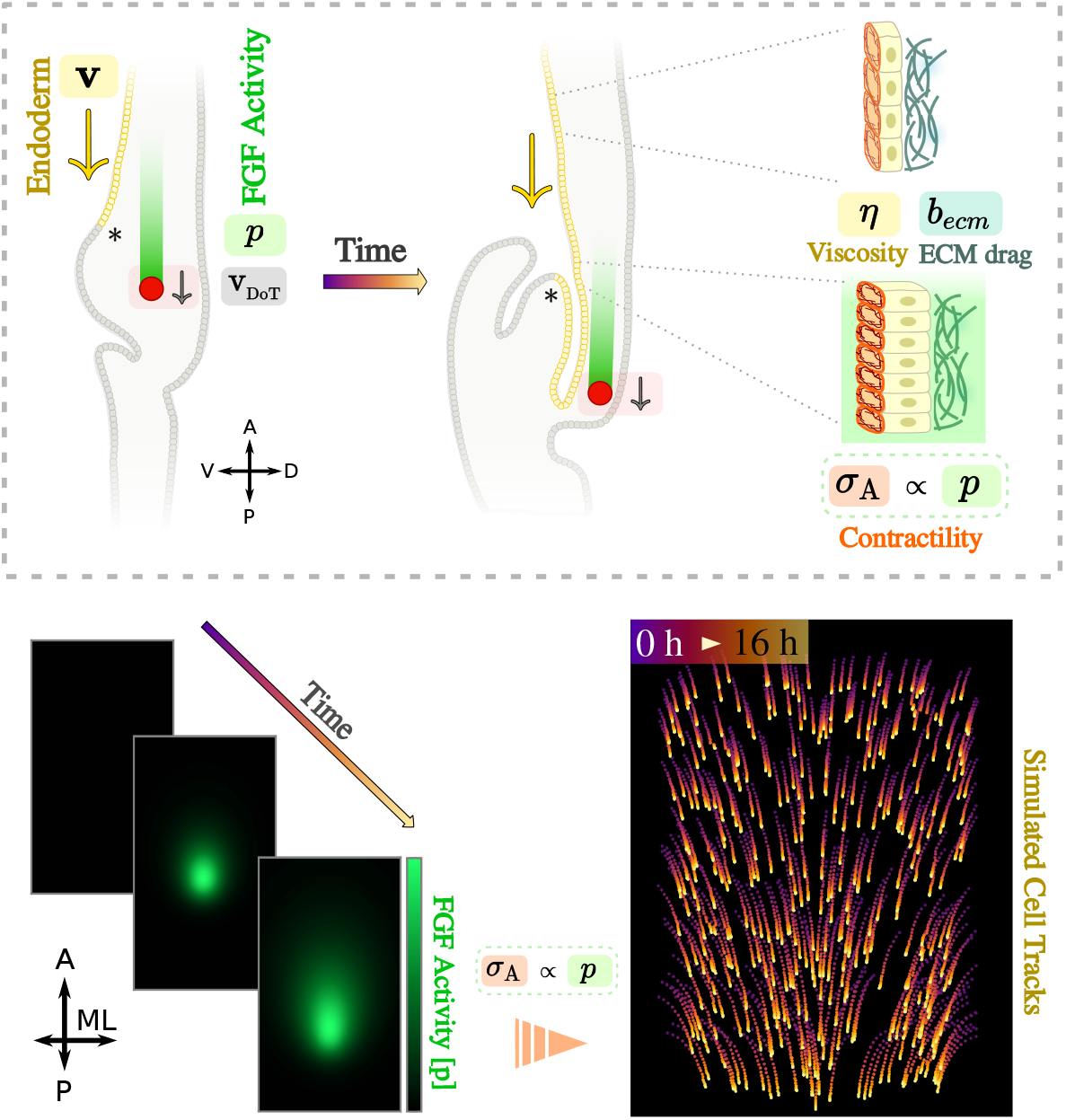

**Summary statement:** This study employs a mathematical model to investigate the interplay between morphogen gradients and tissue mechanics in regulating the collective cell movements that drive hindgut morphogenesis in the chick embryo.

## Introduction

Embryogenesis requires coordination of cell fate specification with profound physical transformations during development. Accordingly, there has been extensive recent work advancing a mechanically-motivated framework for studying morphogenesis [Collinet and Lecuit, 2021, Goodwin and Nelson, 2021, Valet et al., 2022]. These efforts have led to advances in understanding many aspects of development, including how forces deform tissues to create precise morphological outcomes [Heer et al., 2017, Garcia et al., 2019, Nelson and Gleghorn, 2012, Savin et al., 2011, Tallinen et al., 2014], how the subcellular machinery that enacts those forces is regulated [Kasza et al., 2014, Lecuit and Yap, 2015, Martin et al., 2009], and how forces can themselves regulate gene expression [Desprat et al., 2008] and cell behaviors such as proliferation [Godard et al., 2020, Pan et al., 2016], migration [Barriga et al., 2018], and even force generation itself [Lecuit et al., 2011]. Despite these advances, it remains poorly understood – particularly among vertebrates – how forces are patterned during development; how do diffusible signals instruct tissue-scale forces to drive precise morphogenetic events [Lenne et al., 2021]. Indeed, mechanical and biochemical descriptions of vertebrate development have often been invoked as orthogonal to one another, with physical mechanisms such as buckling often viewed as mutually exclusive with those regulated by diffusible signals such as Turing patterns [Durel and Nerurkar, 2020]. As a result, the integration of biochemical cues with a mechanical description of morphogenesis has received less attention. Chemo-mechanical coupling has received recent attention in relation to how tissue flows redistribute the acto-myosin machinery that drive them [Bailles et al., 2022, Serra et al., 2021, Caldarelli et al., 2021, Hannezo et al., 2015]. Moreover, a handful of recent studies have also investigated the role of diffusible signals such as morphogens in regulating tissue-scale mechanics during embryogenesis [Garcia et al., 2019, Pinheiro et al., 2022] and pattern formation *in vitro* [Fei et al., 2020, Boocock et al., 2023]. Nonetheless, it remains unclear how physical properties of embryonic tissue are integrated with transport properties of morphogenic signals to guide vertebrate embryogenesis. The present work employs a mathematical model to study these links during morphogenesis of the avian hindgut.

The gut tube is an endodermally derived epithelial structure in the early embryo that gives rise to the inner lining of the entire respiratory and gastrointestinal tracts [Zorn and Wells, 2009]. During gastrulation in amniotes (birds and mammals), endodermal progenitors ingress from the epiblast through the primitive streak and re-epithelialize on the ventral surface of the embryo. How this epithelial sheet is then internalized and transformed into a tube remains poorly understood, despite the fundamental importance for this process in establishing the body plan. While ultimately forming a single continuous tube along the antero-posterior extent of the embryo, the three segments of the gut tube, foregut, midgut, and hindgut, initiate at distinct developmental stages, forming by seemingly disparate physical and molecular mechanisms [Bellairs and Osmond, 2014, Durel and Nerurkar, 2020, Miller et al., 1999, Hosseini et al., 2017, Nerurkar et al., 2019]. The gut tube begins forming at the anterior end of the embryo with the initiation of a crescent shaped fold, the anterior intestinal portal (AIP), which is then displaced posteriorly, pulling the endoderm ventrally across itself to elongate into a blind-ended tube that will form the foregut. Subsequently, at the posterior end of the embryo, a second invagination forms, termed the caudal intestinal portal (CIP). However, unlike the AIP in foregut morphogenesis, here the CIP remains stationary, and instead it is the movement of endoderm cells through the CIP into the tailbud that drives elongation of the forming hindgut [Nerurkar et al., 2019]. Subsequently, the endoderm between the AIP and CIP folds medially to enclose the presumptive midgut [Miller et al., 1999].

During hindgut morphogenesis, endoderm cells move collectively through the CIP, outpacing outgrowth of the tailbud [Nerurkar et al., 2019]. This mismatch in elongation between the endoderm and tailbud is accommodated by dorso-ventral inversion of the epithelial sheet, leading to progressive folding and elongation of the forming hindgut tube (Fig 1A, B). As such, these collective cell movements are a central driver of hindgut formation, and it is critically important that these movements be coordinated with posterior elongation of the embryo. This is achieved by dependence of both events on the same molecular cue: a long-range gradient of Fibroblast Growth Factor (FGF) ligands [Bénazéraf et al., 2010, Dubrulle and Pourquié, 2004, Nerurkar et al., 2019]. *Fgf4* and *Fgf8* are expressed in a posterior-to-anterior gradient in the tailbud mesenchyme and presomitic mesoderm [Boulet and Capecchi, 2012, Dubrulle et al., 2001, Crossley and Martin, 1995], tissues that lie immediately dorsal to the endoderm. Endoderm cells convert this gradient in FGF ligands into a proportional gradient of acto-myosin activity, resulting in a force imbalance that pulls cells toward the tail end of the embryo (Fig 1C) [Nerurkar et al., 2019]. This leads to a positive feedback loop whereby anterior cells exposed to low FGF levels are displaced passively to higher FGF levels as they are pulled posteriorly by contracting cells at higher FGF concentrations near the CIP, resulting in a progressive recruitment of cells toward the posterior end of the forming hindgut. This is counteracted by movement of the FGF gradient itself, which is displaced posteriorly as the embryo elongates [Boulet and Capecchi, 2012, Bénazéraf et al., 2010, Sawada et al., 2001, Dubrulle et al., 2001].

**Figure 1.**
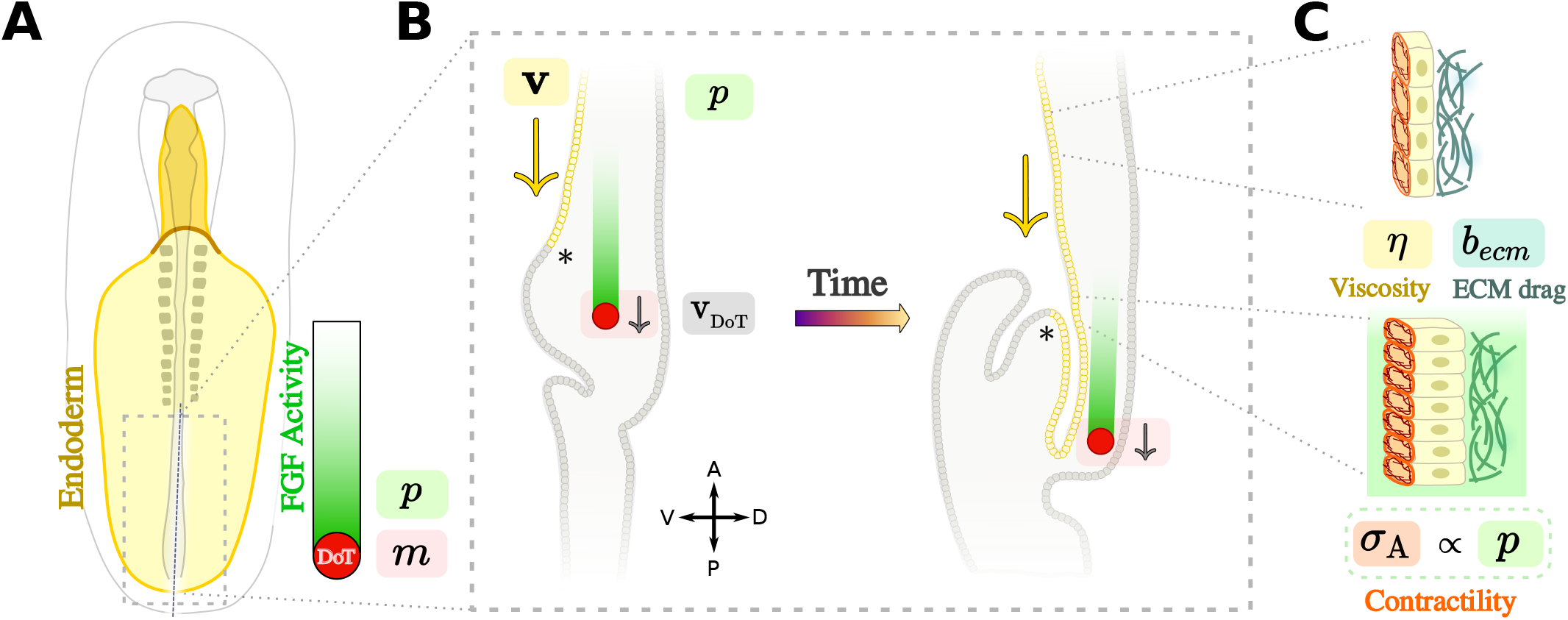
Schematic overview of the signaling and biophysical aspects of avian hindgut morphogenesis. (**A**) A chick embryo (Hamburger Hamilton stage 10, approx. 1.5 days of development) with the endodermal epithelium (ventral) shown in yellow. At this stage, an antero-posterior gradient of FGF protein (green) has already been established, owing to cells in the presomitic mesoderm that are shed from the domain of active transcription (DoT, red) where self-renewing cells actively transcribe *Fgf*. (**B**) Sagittal view of the tailbud region illustrating hindgut tube formation. The endodermal sheet (yellow) and the DoT move at different rates (**v** vs v_DoT_). (**C**) The mechanical properties of the endoderm affect its subsequent movements via activation of isotropic contraction as a result of FGF signalling. (Anatomical axes: A - Anterior, P - Posterior, V - Ventral, D - Dorsal). Asterisk (*) in (B) marks the boundary between gut-forming and extra-embryonic endoderm. *m* is the *Fgf* mRNA concentration, *p* the FGF protein concentration, *η* the endodermal epithelium viscosity, *b*_*ecm*_ the viscous drag force coefficient from ECM interactions, and *σ*_*A*_ the active stress from cell acto-myosin contractility.

Classically, morphogen gradients have been thought to arise due to diffusion of protein from a localized source across a field of cells [Driever and Nüsslein-Volhard, 1988, Yu et al., 2009, Crick, 1970]. However, the posterior FGF gradient instead forms by posterior-ward displacement of a restricted population of self-renewing cells that actively transcribe *Fgf8* ; cells are shed from this domain as axis elongation proceeds, retaining unstable transcripts that are degraded over time [Dubrulle and Pourquié, 2004]. As a result, anterior (“older”) cells retain fewer transcripts than posterior (“newer”) cells. When coupled with translation and diffusion of the resulting FGF protein, this results in the formation of an anisotropic gradient of FGF protein [Dubrulle and Pourquié, 2004] that is accentuated along the antero-posterior axis. Mathematical models of somitogenesis [Baker et al., 2008] have captured the behavior of one-dimensional protein gradient formation through this mechanism, providing important insights into the underlying cell behaviors and regulatory networks that balance the persistence of a stable a domain of active transcription with axis elongation [Harrison et al., 2011], as well as dynamic consequences of the protein gradient for somitogenesis [Baker et al., 2008]. Because the posterior FGF gradient has largely been studied in the context of presomitic mesoderm and axis-elongation, less attention has been paid to the two-dimensional aspects of gradient formation. However, in the hindgut, where directional cell movements are thought to arise due to anisotropy of the gradient, it is necessary to consider how two-dimensional aspects of the FGF gradient inform the mechanics of cell movement in the endoderm.

The present study formulates a 2-D chemo-mechanical mathematical model to investigate how tissue mechanics and spatial control of cell behavior by diffusible signals coordinately regulate cell movements in the hindgut endoderm. We first constructed a 2-D model of FGF gradient formation to investigate how transport properties (e.g., production, stability, and diffusion) of FGF proteins and mRNA interact with the rate of axis elongation to control the shape and extent of the FGF gradient. Next, we developed a continuum model of the endoderm as an active fluid, examining how FGF concentration-dependent active tension regulates tissue flows during extension of the hindgut. The model replicates several experimentally observed behaviors of the posterior chick endoderm, and provides evidence that graded, but isotropic (equal in all directions) cell contractility can drive highly anisotropic (directionally biased) collective movements in the absence of any other asymmetries. Finally, exploring the parameter space of the model provides new insight into the efficiency and evolvability of morphogenic mechanisms, to provide a greater understanding of how chemo-mechanical coupling across the mesoderm and endoderm coordinates hindgut elongation with outgrowth of the tailbud.

## Materials and Methods

### Quantification of FGF activity gradient in the chick endoderm

Fertilized White Leghorn chicken eggs were incubated in a humidified, temperature-controlled chamber (60%, 38° C). Embryos were harvested on filter paper rings [Chapman et al., 2001] at Hamburger-Hamilton (HH) stages 9 and 14, and endoderm-specific electroporation carried out as described previously [Nerurkar et al., 2019]. Briefly, 5*μ*g/*μ*L of plasmid DNA in 5% sucrose and 0.1% Fast Green was injected onto the ventral surface of explanted embryos in a PBS bath, followed by electroporation with the NEPA 21 transfection system (Nepa Gene), using a pair of square electrodes with the following pulse parameters: three 40V poring pulses of 0.1 msec duration, separated by 50 msec, with 10% decay between each pulse, followed by five 4V transfer pulses of 5 msec duration, separated by 50 msec, with 40% decay between each pulse [Nerurkar et al., 2019]. Downstream activity of the FGF pathway was visualized in the posterior endoderm by electroporation of a reporter plasmid consisting of the mouse DUSP6 promoter driving expression of either nuclear-localized eGFP or mScarlet [Nerurkar et al., 2019, Ekerot et al., 2008]. The DUSP6 reporter was co-electroporated with a ubiquitous reporter driving expression of BFP to serve as an electroporation control, and FGF activity was quantified by normalizing the DUSP6 driven signal intensity to that of the ubiquitous reporter [Nerurkar et al., 2019, Wang et al., 2014]. Electroporated embryos were imaged on an AxioZoom v1.2 macroscope (Zeiss) after 6 hours of incubation. Post-processing of images was carried out using the Napari viewer [Sofroniew et al., 2022], Napari Assistant plugin [Haase et al., 2023] and sci-kit image library [van der Walt et al., 2014]. Briefly, Otsu’s method [Otsu, 1979] was used to perform automatic image thresholding and detect DUSP6-driven fluorescence in cells, and maximal Dusp6 intensity values per nucleus were normalized by maximal BFP (control) intensity. The entire image was then binned into a 38 × 10 grid { antero-posterior x medio-lateral }, and normalized DUSP6 intensity values were averaged per bin.

### Two-dimensional model of FGF gradient formation

Previous work by Harrison and colleagues [Harrison et al., 2011] established a one dimensional model of FGF gradient formation that relates posterior displacement of a restricted population of cells actively transcribing *Fgf8* mRNA, transcript decay, and diffusion of translated FGF8 protein to establish a long-range gradient of FGF ligand. However, in order to ask how establishment and shape of the FGF protein gradient influences cell movement anisotropy in the neighboring endoderm, we first generalized this model into two dimensions. A detailed description of the model formulation and solution is provided in Supp. Methods, with key aspects of the model outlined below. To construct a 2-D model of FGF gradient formation in the posterior embryo, we modeled transport of two species, *Fgf8* mRNA (*m*) and the FGF8 protein (*p*). Production of *m* was restricted to a posterior region, the Domain of Transcription (DoT), that moves posteriorly with velocity v_DoT_ (Fig 1A,B). Transcripts left in the wake of the DoT as it progresses posteriorly were specified to degrade in proportion to transcript concentration *m*. FGF protein *p* was modeled as having a rate of production proportional to the concentration of *m* to signify translation from mRNA, and a degradation rate proportional to protein concentration *p*. Finally, while mRNA remains restricted within cells, FGF protein *p* was permitted to diffuse. When formulated in a reference frame that moves with the DoT, the associated mass balance results in a pair of coupled partial differential equations:

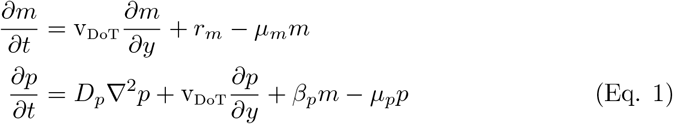

where *m* is the concentration of *Fgf* mRNA, v_DoT_ is the velocity with which the embryo elongates posteriorly (hence the velocity of the DoT), *r*_*m*_ is the rate of mRNA synthesis within the DoT, *μ*_*m*_ is rate of mRNA degradation, *p* is the concentration of FGF protein, *D*_*p*_ is the diffusion coefficient of FGF protein, *β*_*p*_ is the rate of protein production from translation, and *μ*_*p*_ is the rate of protein degradation. Advection terms (containing v_DoT_) are included in Eq. 1 to account for the moving frame of reference in which balance equations were derived. v_DoT_ was assumed to be constant, based on previous studies indicating that axis elongation rate in the chick embryo is approximately constant at ∼50*μ*m/hr through the stages of interest [Xiong et al., 2020, Bénazéraf et al., 2010]. The equations were solved on a rectangular, 2-D domain approximating the relatively flat posterior endoderm of an HH 15 chick embryo. However, to limit the influence of artificially simple boundary conditions on the overall behavior of the model, the domain size was chosen to be vastly larger than the length-scale of the embryo proper: 20 mm × 60 mm corresponding to the medio-lateral and antero-posterior axes, respectively.

The model was de-dimensionalized by defining scaled variables:

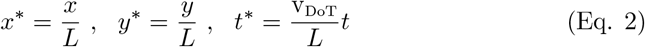

where *L* approximately represents the characteristic length-scale of the FGF gradient along the antero-posterior axis. Substituting Eq. 2 into Eq. 1 produces the dimensionless form:

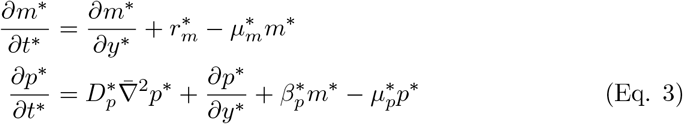

where the six model parameters have been reduced to four dimensionless ones:

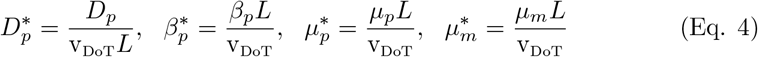

the asterisk denotes a unit-less parameter, and 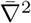 corresponds to the Laplacian with respect to the scaled spatial coordinates of Eq. 2.

Eq. 3 was solved on a rectangular domain representing half of the embryo to account for bilateral symmetry of the FGF gradient [Dubrulle et al., 2001, Dubrulle and Pourquié, 2004] and endoderm cell movements [Nerurkar et al., 2019]. Accordingly, the left boundary of the domain was subject to a symmetry boundary condition. The remaining three boundaries - anterior, lateral, and posterior to the DoT – were assumed to be sufficiently far away that protein and mRNA concentration were zero. In addition, a finer mesh density was applied within a 2.5 mm x 5 mm subdomain corresponding to the gut-forming endoderm. It was assumed that initially, mRNA and protein concentrations are zero throughout the domain. The pair of differential equations describing mRNA and protein distributions subject to these boundary and initial conditions were solved numerically using the open-source computing platform FEniCS [Logg and Wells, 2010, Logg et al., 2012, A. Logg et al., 2012, Kirby, 2004, Kirby and Logg, 2006, R. C. Kirby, 2012] to obtain time-varying spatial distributions of Fgf8 mRNA and FGF8 protein for a given set of parameter values (below).

The value of protein kinetic parameters were determined from FGF activity measured experimentally using the FGF-responsive DUSP6 reporter. For simplicity, we performed curve fits of an analytical solution to the 1-D equivalent of Eq. 3, derived previously by Harrison and colleagues [Harrison et al., 2011] (see Supp. Methods), to DUSP6 reporter activity at HH stage 10 (n=9) and HH 15 (n=10) to simultaneously obtain values for *D*_*p*_, *μ*_*p*_ and *β*_*p*_. In order to examine how each model parameter influences overall protein distribution in the model, four metrics were used (Fig 4A). The first is a localized measure, maximal protein concentration 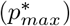. Next, to describe the 2-D shape of the gradient, iso-concentration-lines at 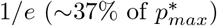 of of the maximum concentration were isolated and shapes were quantified by an overall antero-posterior gradient length 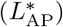, ellipticity (*T*), and asymmetry (*λ*), based on approximation of the shape as an ovoid (Fig 4A, Supp. Methods). The time dependent transport equations for the protein concentration were run until a steady state solution was achieved, which was determined to have been reached after at least four successive time steps show a less than 0.5% difference from the previous time point (SFig 1).

The effects of each parameter on model behavior were determined by sweeping its value while holding all other parameters at ‘baseline/physiologic’ values (as measured by experiment or from literature). The upper and lower bounds for the values of each swept parameter were chosen to be 1-3 orders of magnitude away from the baseline value, with five or more values per decade-interval sampled. For more details on the swept values of each parameter see Table 1.

**Table 1.** Baseline values used for dimensionless parameters in simulations (and estimated experimentally for 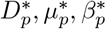). 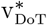 and *η*^*^ represent scaled parameters normalized to experimentally measured axis elongation rate and an arbitrary reference viscosity, respectively. ^†^Ranges (for parametric sweeps) are reported in log-10-space as [start, stop, steps].

| Parameter | Value | Range<sup>†</sup> |
| --- | --- | --- |
| **FGF Transport** |  |  |
| $D_p^*$ | 0.18 | [-1,2,25] |
| $\mu_p^*$ | 1.5 | [-1,2,25] |
| $\beta_p^*$ | 5 | [-1,2,25] |
| $\mu_m^*$ | 20 | [0,3,25] |
| $v_{DoT}^*$ | 1 | [-1,1,31] |
| **Tissue Mechanics** |  |  |
| $\eta^*$ | 1 | [-1,2,25] |
| $K^*$ | 10 | [-1,2,25] |
| $b_{ecm}^*$ | 0.1 | [-2,1,25] |
| $\alpha^*$ | 10 | [-1,2,25] |

### 2-D model of collective movements driving hindgut formation

Upon solution of the 2-D model of FGF gradient formation described above, we next constructed a model of cell movements in the endoderm that occur in response to this gradient [Nerurkar et al., 2019]. The tissue is modeled as a thin two dimensional active fluid, where FGF-mediated acto-myosin contractility induces a multi-cellular flow. Accordingly, total stress (*σ*_*T*_) in the endoderm was decomposed into the sum of a passive (*σ*_*P*_) and active (*σ*_*A*_) stress:

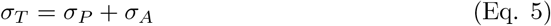

The constitutive law describing passive mechanical properties of the endoderm was assumed to take the form of a viscous compressible fluid such that

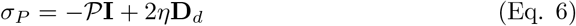

where 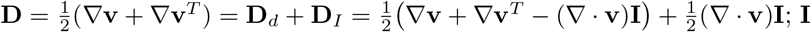 is the identity tensor, **v** = [*v*_*x*_, *v*_*y*_] is the velocity, 𝒫 is the pressure, and *η* the viscosity.

The active stress, representing acto-myosin activity of endoderm cells generated in response to FGF8 (Fig 1C, was assumed for simplicity to vary in proportion to FGF protein concentration

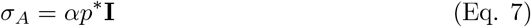

where *α* is a factor to convert from units of concentration to units of stress and *p*^*^ is the FGF protein concentration determined by solving Eq. 3 above. Owing to the symmetric cell shape of endoderm cells subject to high FGF8 in the posterior endoderm [Nerurkar et al., 2019], we have assumed that contraction is isotropic within the epithelial plane. The movement of the endoderm was assumed to be passively resisted by extracellular matrix. Because large extracellular matrix displacements have been observed in several contexts of avian morphogenesis [Aleksandrova et al., 2015, Zamir et al., 2008, Bénazéraf et al., 2010], we assumed that the matrix acts to resist cell movements as a viscous drag rather than by elastic deformation. Accordingly, in the limit of slow flows associated with morphogenesis, we can neglect inertia so that local force balance reads:

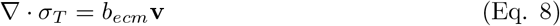

where *b*_*ecm*_ is the viscous drag coefficient resulting from ECM interactions between endoderm cells and their basement membrane. While one could envision that in three dimensions, the endoderm behaves similar to an incompressible fluid on short time scales (meaning all tissue deformations must be volume preserving when proliferation is ignored), in two dimensions it is necessary to accommodate area changes associated with cell contractility, and so the continuity equation was taken to be of the form [Serra et al., 2021]

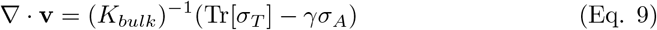

where *K*_*bulk*_ is the bulk viscosity and *γ* is a factor that controls the amount of out of plane motion accounted for by the active stress. Substituting equations Eq. 5, Eq. 6, and Eq. 7 into Eq. 9 provides an explicit relation for pressure

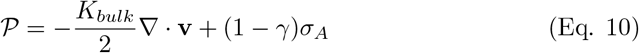

such that the only independent variable remaining is the velocity field. Combining equations Eq. 5, Eq. 8, and Eq. 10 reduces the force balance to:

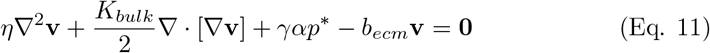

which can be simplified upon de-dimensionalization to:

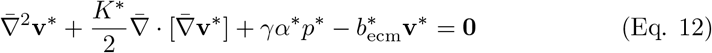

where

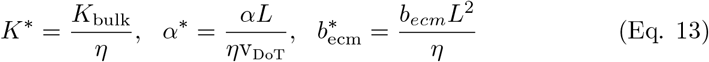

are the dimensionless parameters describing tissue mechanics, and spatial and time variables are scaled as in Eq. 2. As with Eq. 3 describing FGF protein concentration, Eq. 12 was solved on a rectangular domain representing half of the bilaterally symmetric embryo, such that a symmetry condition was applied to the medial (left) boundary, while the anterior, lateral, and posterior boundaries of the domain were assumed to be sufficiently distant from the region of interest that **v** = 0 was applied at each. The steady state solution of the FGF transport equations, specifically the scalar field of the protein concentration, *p*^*^, was passed to the tissue mechanics equations as an input field for defining the active stress Eq. 7. Transport and mechanics equations were solved on the same finite element mesh in the moving frame of reference. To compare simulation results across varied input parameters, the following output metrics were defined: (1) *Q*_AP_ - Bulk Tissue flow in the antero-posterior axis, defined as the y-axis component of the velocity field integrated over the entire domain anterior to the maximal *p*^*^ concentration (to capture the full capacity of the solved velocity field to accommodate morphogenetic movements) (2) *v*_*max*_ - maximum posterior-ward velocity (reported as an absolute value), and (3) *Q*_AP_*/Q*_ML_ - tissue flow ratio, defined as the ratio of the antero-posterior to medio-lateral bulk tissue flow.

### Time lapse imaging and simulated cell tracks

Live imaging of endoderm cell movements was performed as described previously [Nerurkar et al., 2019]. In brief, embryos electroporated with pCAG-H2B-GFP, which drives nuclear-localized GFP expression under control of a ubiquitous promoter, were imaged in a Zeiss LSM confocal microscope equipped with two-photon excitation for 16 hours at 37 ° C. Resulting cell tracks were visualized by depth-coded projection of the time series in Fiji, and compared with cell track simulations from the solved velocity field of Eq. 12, using `scipy.interpolate.interp2d` for the spatial interpolation, and the forward Euler method for time integration.

## Results

### 2-D mRNA decay model quantitatively replicates embryonic FGF activity gradient in the endoderm

In order to study 2-D endoderm cell movements in terms of tissue mechanics and transport properties of FGF ligands, we first formulated a 2-D model describing the establishment and maintenance of an anisotropic, two-dimensional gradient of FGF protein. The long-range FGF gradient in the posterior embryo arises due to progressive posterior displacement of a population of cells actively transcribing *Fgf8*, coupled with mRNA decay and diffusion of translated FGF protein [Dubrulle and Pourquié, 2004]. We formulated a 2-D reaction-diffusion-advection framework of coupled partial differential equations that builds on the 1-D model previously developed by Harrison and colleagues [Harrison et al., 2011]. It was assumed that the region of cells actively transcribing *Fgf8* (the domain of transcription, DoT) moves posteriorly with a velocity 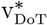 as the embryonic body elongates and the tailbud moves posteriorly (Fig 1). This leaves in its wake a trail of mRNA, which due to a constant rate of mRNA decay results in a gradient of mRNA concentration *m*^*^ that is progressively displaced posteriorly (Fig 2A, Supplemental Movie 1). Posterior-ward movement of the domain of active transcription also introduces anisotropy to the mRNA gradient, which becomes progressively elongated along the antero-posterior axis. The balance equation describing mRNA transport was then coupled to a second partial differential equation describing protein distribution *p*^*^. FGF protein was assumed to be produced by translation at a rate proportional to mRNA concentration, degraded in proportion to protein concentration, and unlike mRNA, was free to diffuse across the tissue. This resulted in a traveling, long-range gradient of FGF protein that extends beyond the mRNA gradient, and has a more pronounced anisotropy (Fig 2). When solved in a frame of reference traveling with the domain of transcription, the model smoothly approaches a stable steady state solution over time (Fig 2B).

**Figure 2.**
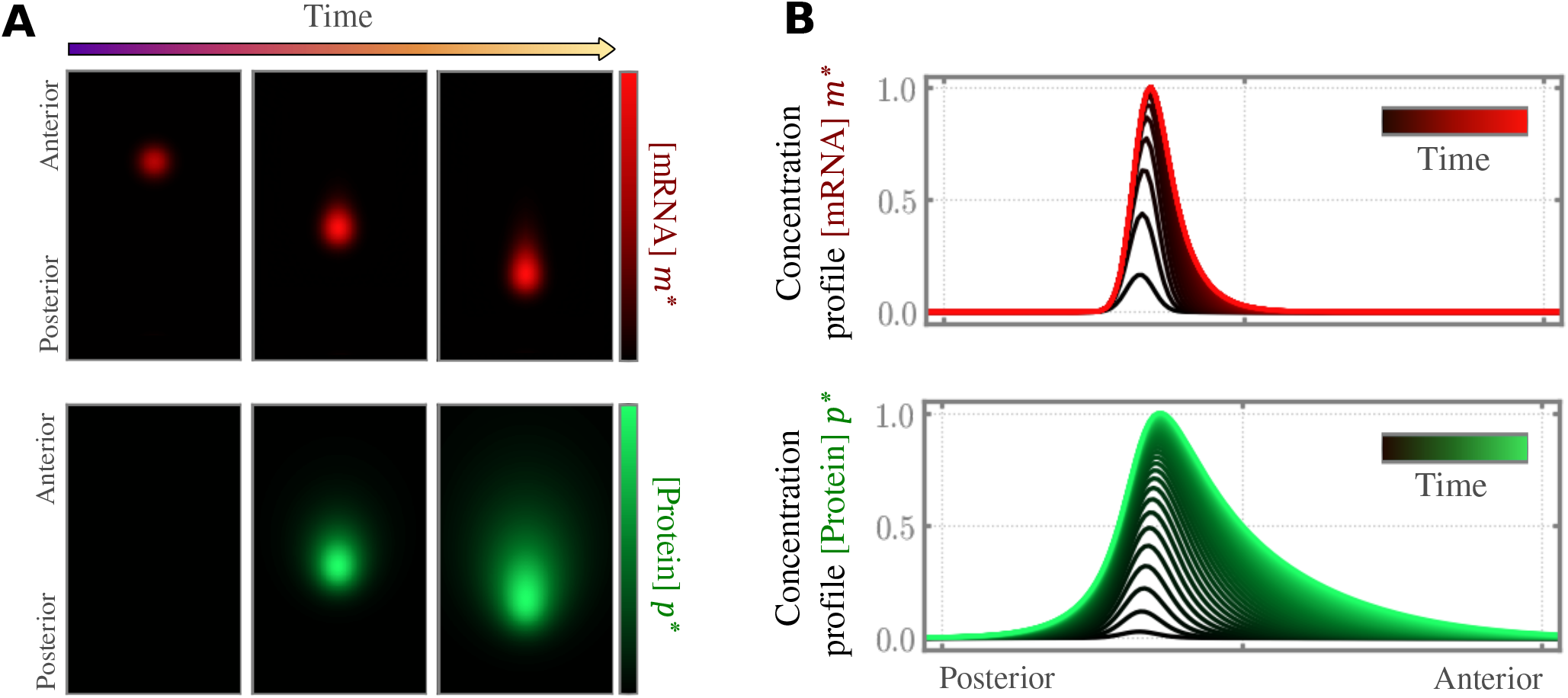
Simulation of the mRNA decay model and corresponding mRNA and protein concentration profiles. (**A**) 2-D concentration profiles of mRNA (top, red) and protein (bottom, green) as visualized in the fixed/laboratory frame of reference as the embryo elongates and the Domain of active Transcription (DoT) is displaced posteriorly. (**B**) The temporal evolution of the concentration profiles for mRNA and protein along the midline of the embryo (now in the moving frame of reference. Line color indicates temporal progression in the concentration profile from the initial time point (black) to the steady state mRNA (red) and protein (green) concentrations; the time step is constant between each successive curve.

We next tested experimentally whether, as predicted by the model, the FGF gradient does in fact stabilize during axis elongation in the chick embryo. To do so, FGF activity in the endoderm was quantified by electroporation of a plasmid encoding a nuclear-localized fluorescent protein downstream of the FGF-responsive DUSP6 promoter; signal was normalized to that from a co-electroporated ubiquitous reporter serving as an electroporation control (Fig 3A [Nerurkar et al., 2019]). The antero-posterior shape of the FGF activity gradient was compared along the embryonic midline at HH stage 10, prior to the onset of hindgut formation, and HH 15, when hindgut formation is already underway. We reasoned that a similar distribution of FGF activity in the endoderm across these two stages would indicate that the gradient is stably established prior to the onset of collective cell movements driving hindgut formation, and as a result, that the steady state solution could be used in all subsequent studies on downstream mechanics of endoderm flows. Qualitatively, no difference in the antero-posterior profile of the FGF activity gradient was observed between HH 10 and HH 15 (Fig 3B). To quantitatively test this, DUSP6 reporter activity was quantified in terms of model parameters describing FGF protein transport: diffusion 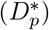, protein decay rate 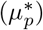 and protein translation rate (*β*_*p*_). In brief, a previously derived analytical solution to the 1-D equivalent of the FGF transport model described above [Harrison et al., 2011] was fit to these experimental data (dashed lines, Fig 3B) using an initial guess of the mRNA decay parameter *μ*_*m*_, which was then revised via 2-D correlation to experimental results (Fig 3D, E). When DUSP6 reporter activity was quantified in terms of model parameters, indeed we observed no difference diffusion 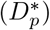, protein decay rate 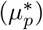 and protein translation rate (*β*_*p*_) (Fig 3C) between stages HH10 and 15. These results further validate the model prediction that gradient formation approaches a steady state solution prior to hindgut formation, and that axis elongation results in a posterior shift in the FGF gradient without changing its shape. In addition, the quantification of model parameter values from fitting experimental data provides an important physiologic baseline from which parametric sweeps can be conducted to understand how each parameter influences the dynamics and shape of FGF gradient establishment in 2D. Therefore, the model captures the basic shape and extent of the FGF activity gradient observed in vivo using model parameters that are constrained by experiment.

**Figure 3.**
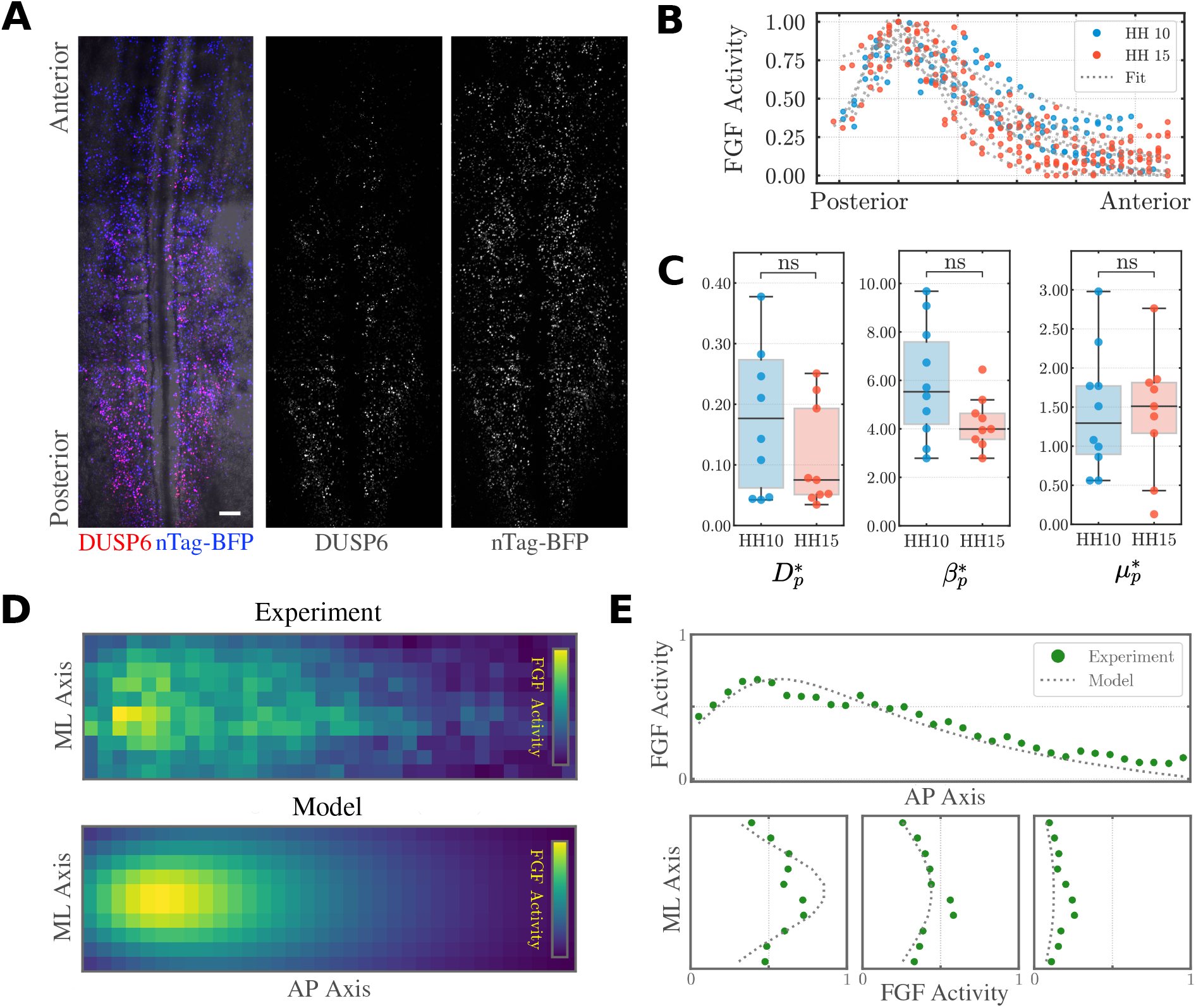
Measurement of FGF pathway activity in vivo. (**A**) Representative ventral view of stage HH 15 chick embryo following co-electroporation of posterior endoderm with the FGF-responsive DUSP6-mScarlet reporter and the ubiquitous reporter nTag-BFP (control). Scale Bar 100*μ*m. (**B**) DUSP6 reporter intensity normalized to nTag-BFP and plotted as a medio-lateral average as a function of antero-posterior position at stages HH 10 (blue) and HH 15 (red); dashed lines indicate model fit to experimental data used to establish physiologic values of model parameters 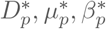 (**C**) FGF transport parameters obtained from fitting DUSP6 data at stages HH 10 (n=9) and HH 15 (n=10); n.s. = no significant difference as established by two-sided Mann-Whitney-Wilcoxon Test. (**D**) Comparison of 2-D representation of FGF activity from averaged Dusp6-mScarlet/nTag-BFP electroporation and model simulated FGF activity (ML/AP = medio-lateral/antero-posterior axes). (**E**) Comparison of FGF activity on virtual sagittal and transverse slices.

### mRNA and protein kinetics affect morphogen gradient shape

To characterize 2-D FGF concentration profiles, which are the primary output of the FGF transport model, isoline contours corresponding to 1*/e* of the maximal protein concentration (in other words the distance at which protein concentration has decreased to approximately 37% of its peak) were quantified with respect to their ellipticity (*T*) and asymmetry (*λ*) (Fig 4A) [Baker, 2002]. *T* = *λ* = 1 indicates a radially symmetric (isotropic) protein concentration profile one would expect if the gradient arose by isotropic diffusion from a fixed source; *T <* 1 indicates that the gradient is elongated in the antero-posterior direction, while *λ <* 1 indicates asymmetry along the antero-posterior axis that is more posteriorly-biased (Fig 4A). In addition to gradient shape, we quantified the maximum protein concentration 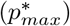 and scaled length of the gradient 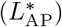.

**Figure 4.**
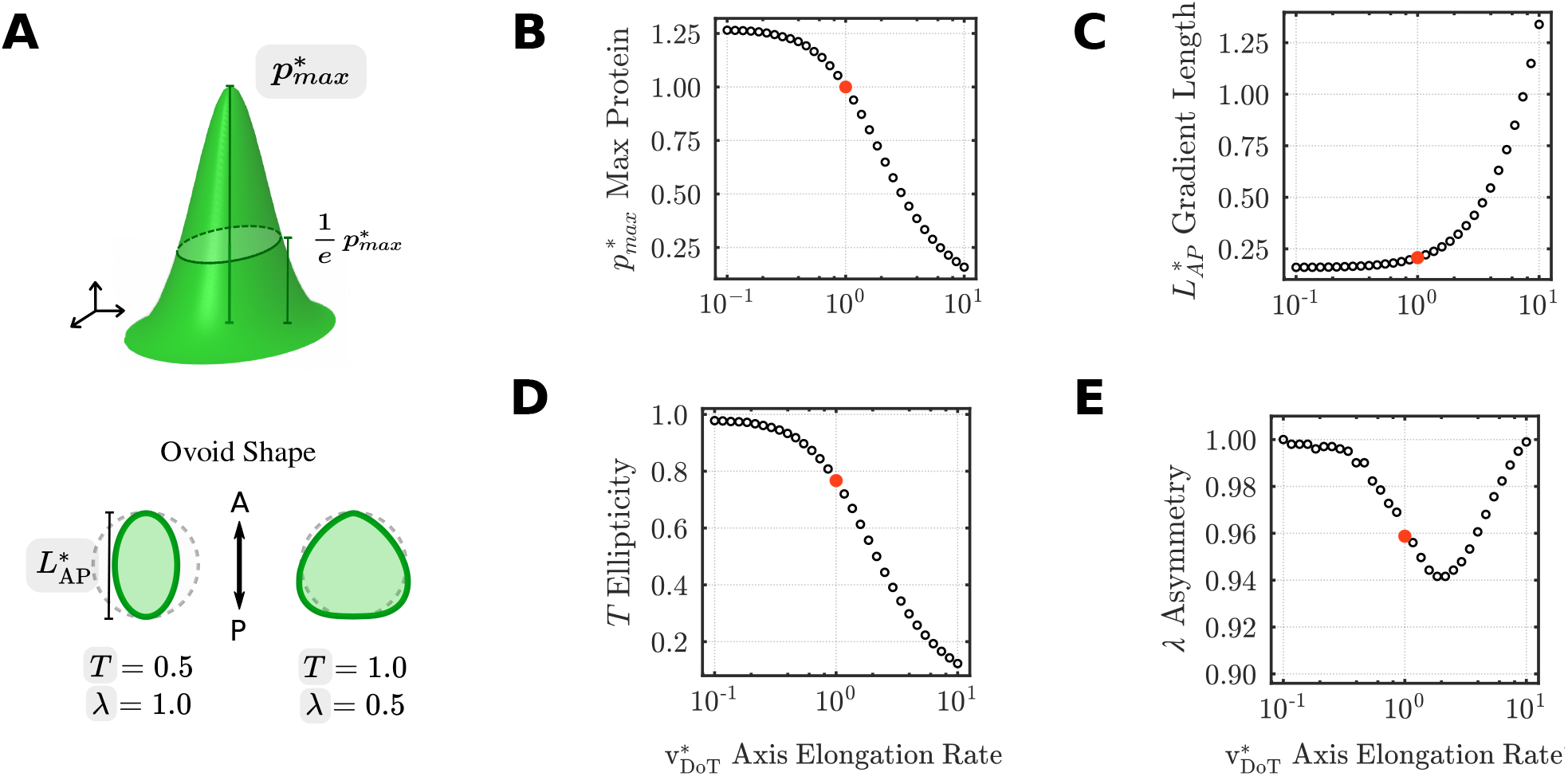
(**A**) Schematic representation of output metrics to characterize gradient shape. (**B-E**) Parametric sweeps on 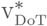; red circle denotes the physiological value of v_DoT_ corresponding to the rate of axis elongation measured experimentally in prior studies (see Methods). Protein levels were normalized to the maximum protein value from the physiological set of parameters.

Given that scaling of the coupled partial differential equations governing FGF gradient formation resulted in absorption of v_DoT_ into the scaled parameters Eq. 4, the effects of changes in this key parameter were first isolated to simplify the interpretation of subsequent parametric sweeps. In these simulations, changes in 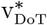 can be interpreted broadly as changes in the rate of embryonic axis elongation, assuming that the domain of cells actively transcribing Fgf ligands are displaced posteriorly with the same rate [Dubrulle and Pourquié, 2004, Bénazéraf et al., 2010]. Increasing 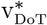 reduced 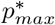, owing to a constant rate of transcription and translation resulting in similar amounts of total FGF protein, but distributed across a broader domain as the embryo elongates posteriorly at higher speeds (Fig 4B). 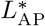, indicating the antero-posterior length over which the FGF gradient acts, varied directly with 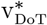 (Fig 4C), confirming the central role of axis elongation in the establishment of the long-range FGF gradient [Harrison et al., 2011, Dubrulle et al., 2001]. This effect was highly anisotropic, as seen by a corresponding decrease in ellipticity (T) (Fig 4D). In other words, elongation of the gradient with faster rates of axis elongation occurred primarily along the antero-posterior axis, with little change in the medio-lateral extent of the gradient (SFig 2). A small, but nonlinear, dependence of asymmetry *λ* on 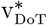 was observed: *λ* changed by less than 10% when 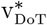 was varied across two orders of magnitude (Fig 4E).

We next considered how the shape of the FGF gradient depends on the scaled mRNA decay rate 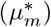 and protein diffusion 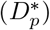. An increase in 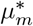 indicates that the system is dominated the instability of FGF transcripts rather than the rate of axis elongation, while an increase in 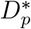 favors protein to diffuse from its site of translation over the effects of axis elongation. Both parameters strongly influence overall features of the FGF gradient (Fig 5A-D). As expected, the maximal protein concentration 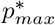 is achieved when both parameters are small (Fig 5A), as this would indicate a long lifetime for FGF transcripts, and that translated protein accumulates locally with limited diffusion from the site of translation. Experimentally-based approximation of 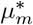 and 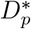 (Fig 3C, Table 1) fall well outside the range of maximal protein concentration, consistent with the importance of the spatial distribution of FGF rather than its absolute levels for axis elongation [Bénazéraf et al., 2010] and hindgut morphogenesis [Nerurkar et al., 2019] (Fig 5A). The antero-posterior extent of the FGF gradient 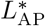 depends nonlinearly on 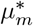, with relatively little change in length as 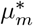 increases beyond baseline values approximated in the chick embryo (Fig 5B). Interestingly, while 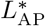 does increase in response to increased diffusion 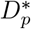, it is far more sensitive to changes in 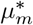 (Fig 5B). Finally, in the limit of increasing 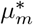 and 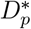, the effects of axis elongation rate were minimized in favor of highly unstable transcripts that are translated into proteins that rapidly diffuse from their source, thereby approaching the case of radially symmetric diffusion from a static source (Fig 5C,D). This further supports the central role of axis elongation in anisotropic gradient formation in the posterior embryo. When comparing the effects of instability at the mRNA and protein level, the maximum protein concentration 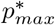 and the antero-posterior length of the protein gradient 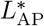 were generally less sensitive to instability at the protein level 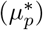 than at the mRNA level across a range of 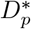 (Fig 5A’,B’). Notably, while protein diffusivity 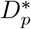 did not have a strong influence on the antero-posterior extent of the FGF gradient across a broad range of mRNA and protein decay rates (Fig 5A’,B’), the ellipticity and asymmetry of the gradient were far more sensitive to 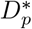 (Fig 5C,D). This suggests that protein diffusivity has a more dominant role in establishing the medio-lateral distribution of FGF proteins than along the antero-posterior axis. The extent to which this influences antero-posterior directed movements in the endoderm is investigated below. Finally, while higher rates of protein production 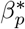 predictably increased the maximum protein concentration 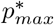, the scaled measures of antero-posterior extent and shape of the FGF gradient were otherwise insensitive to this parameter (SFig 3).

**Figure 5.**
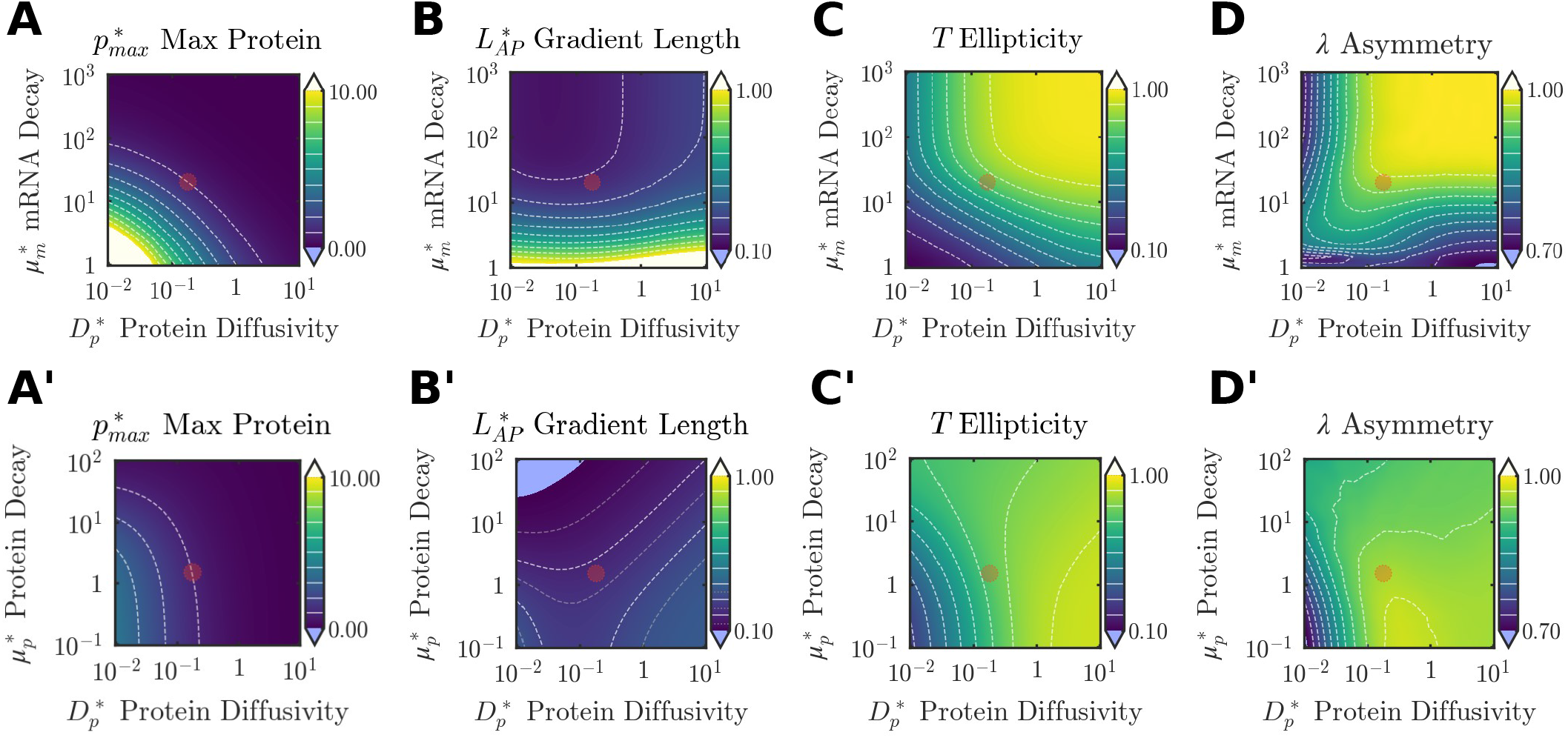
Parametric sweeps on FGF transport properties. Output metrics describing 2D FGF protein gradient shape across parametric sweeps on mRNA decay 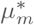 and protein diffusivity 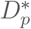 (**A-D**), as well as protein decay 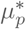 and protein diffusivity 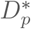 (**A’-D’**). Red circle denotes the baseline values of the swept parameters, as quantified experimentally (Fig 2C). White dashed lines are isolines at 10% intervals of the colorbar range; inn **B’**, grey isolines are at 5% intervals. Protein levels were normalized to the max protein value from the physiological set of parameters.

### Chemomechanical coupling model reproduces key aspects of hindgut morphogenesis

Having formulated a model that describes the 2-D FGF activity gradient in terms of transport properties of FGF transcripts and proteins, we next integrated this with a model aimed at studying endoderm deformations occurring in response to FGF protein concentration. We previously showed that directed, collective cell movements in the posterior avian endoderm arise through conversion of the long-range FGF gradient into a force gradient via RhoA-dependent acto-myosin contractility [Nerurkar et al., 2019]. To conceptualize this in a mathematical model, the endoderm was approximated as an active (force-generating), compressible (capable of volume changes), viscous fluid (mechanical stress is a function of the rate of deformation) with isotropic tensile stress generated in proportion to FGF concentration, wherein FGF concentration was the result of the transport model described above. The non-dimensional parameters describing endoderm tissue mechanics comprise shear viscosity (*η*^*^), bulk/volumetric viscosity (*K*^*^), FGF-responsive contractility (*α*^*^), and viscous effects of the ECM 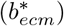, which serve to passively resist cell movements. Unlike transport parameters, which could be approximated directly from experimental measurements of FGF activity, quantification of physical properties was not feasible, and so these parameters have been estimated based on literature (Table 1), and the initial focus is instead on testing qualitative agreement between model and experiments. Indeed, we observed model-simulated cell trajectories that agree well with observed cell movements in the posterior chick embryo during hindgut morphogenesis (Fig 6A, B), and that endoderm movements can outpace axis elongation (*v*_*max*_ *>* 1) as in the chick embryo [Nerurkar et al., 2019]. To test whether the model captures other essential features of the developing hindgut, we examined stress anisotropy accompanying tissue flows. Previous experiments indicated that larger tensile stresses in the endoderm are observed along the medio-lateral axis, rather than along the antero-posterior axis, which is the primary direction of cell movements [Nerurkar et al., 2019]. In agreement with this, the model predicted higher total stresses along the medio-lateral direction (*σ*_ML_) than the antero-posterior direction (*σ*_AP_), despite isotropic contractility driving these tissue movements (Fig 6C). Attenuation of total stress anteriorly was also observed, further supporting experimental findings [Nerurkar et al., 2019].

**Figure 6.**
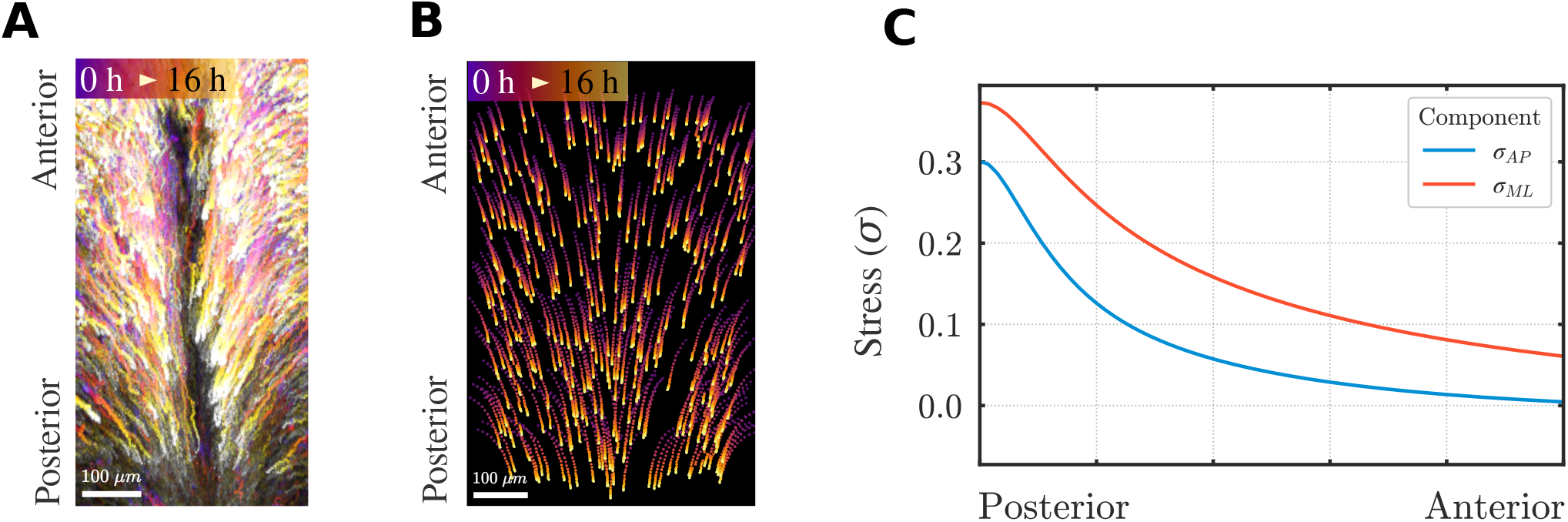
Chemo-mechcanical model replicates key aspecets of avian hindgut morphogenesis. (**A**) Cell tracks from *in vivo* time lapse microscopy of posterior chick endoderm expressing nuclear localized GFP reporter between stages HH 13 and HH18. (**B**) Simulated cell tracks from solution of the chemo-mechanical model of endoderm tissue movements. (**C**) Spatial distribution of relative total stress *σ*_*T*_ (normalized to maximal active stress) along the embryonic midline.

### Isotropic material properties drive directional cell flows in response to FGF activity gradient

Having established a 2-D chemo-mechanical model that replicates basic behaviors observed experimentally during avian hindgut morphogenesis, we next performed parametric analyses to understand how physical properties of the endoderm influence the tissue flows organized by FGF. To simplify characterization of model simulations, three scalar metrics were considered: (1) *Q*_AP_, a bulk antero-posterior tissue flow rate providing a macroscopic readout of overall tissue movements, (2) *v*_*max*_, the maximum (posteriorward) antero-posterior speed, which provides a local measure of tissue movement, and (3) *Q*_AP_/*Q*_ML_, which quantifies the degree of directionality in cell movements by comparing antero-posterior flows to medio-lateral flows.

We first examined how these metrics are altered when FGF transport properties are held fixed at physiologic values (Table 1), and parameters describing endoderm mechanics are varied. The ECM was modeled as a viscous drag force acting to resist cell movements, defined by a linear drag coefficient 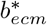. A second, intrinsic resistance to cell movements was provided by the viscous deformability of the endoderm itself, described by *K*^*^. Both parameters had similar effects, with increases in each diminishing the influence of the other, and increasing either 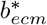 or *K*^*^ reducing *Q*_AP_ and *v*_*max*_ (Fig 7A, B). In other words, at sufficiently high 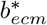, the model was largely insensitive to changes in material properties of the endoderm. In contrast, *K*^*^ and 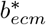 had dissimilar effects on the directionality of tissue flow: increasing *K*^*^ progressively biases the flow toward antero-posterior movement while changes in 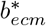 have relatively little effect (Fig 7C).

**Figure 7.**
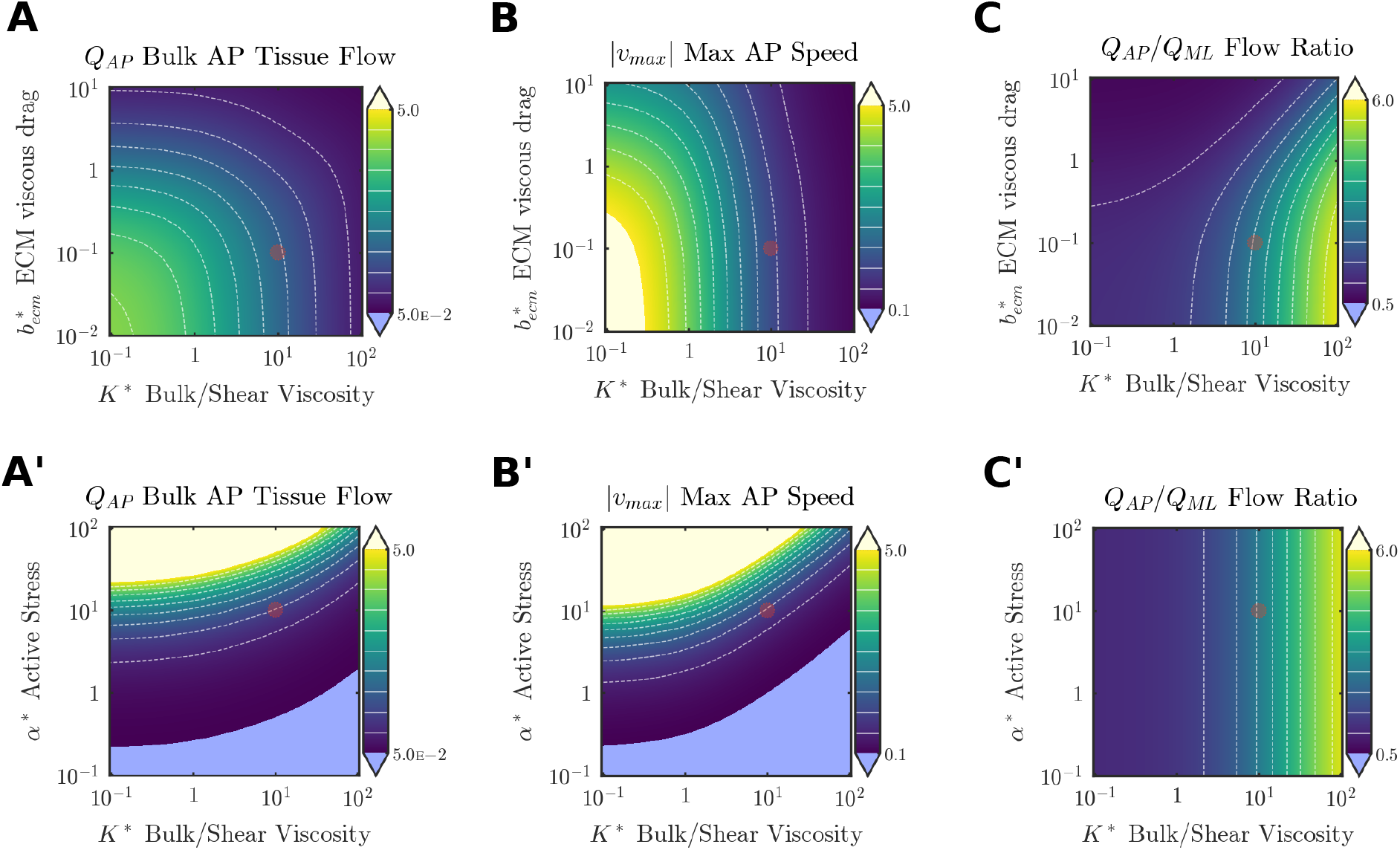
Effects of tissue mechanics parameters on endoderm deformations. (**A-C**) Sweeps of viscous drag from ECM 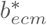 and bulk to shear viscosity ratio *K*^*^. (**A’-C’**) Sweeps of active stress *α*^*^ and bulk to shear viscosity ratio *K*^*^. Red circle indicates baseline parameters. White dashed lines are isolines at 10% intervals of the colorbar range.

We next considered how the responsiveness of endoderm to FGF in terms of actomyosin contractility, described by *α*^*^, influences endoderm deformations. Bulk tissue flows and local maximum velocity increased with *α* *, confirming that increasing the conversion of FGF concentration to acto-myosin activity will coordinately produce larger deformations (Fig 7A’, B’). Antero-posterior tissue flows were only strongly sensitive to material properties of the endoderm at high contractility, with little dependence on *K*^*^ throughout much of the range of *α*^*^ considered (Fig 7A’); as a more local measure of tissue movement, the maximum antero-posterior velocity was somewhat more responsive to changes in *K*^*^ (Fig 7B’). In contrast, however, *K*^*^ strongly influenced the anisotropy of endoderm flow, with higher deformability enhancing the directionality of deformations along the antero-posterior axis (Fig 7C’). Notably, anisotropy was insensitive to contractility *α*^*^ (Fig 7C’), highlighting that anisotropy of the movements stems from the spatial distribution of the active tension gradient, not the magnitude of active tension, which is generated isotropically.

### Axis elongation and mRNA decay strongly influence endoderm deformations driving hindgut morphogenesis

Having analyzed the dependence of the chemo-mechanical model on mechanical parameters of endoderm for a given ‘physiologic’ distribution of FGF ligands, we finally examined interactions between transport properties of FGF and mechanics of the endoderm. De-dimensionalization of the transport and mechanics equations led to absorption of two key properties, v_DoT_ and *η*, respectively, into the remaining model parameters. In other words, all other metrics of transport and mechanics only influence the behavior of the system via their relative magnitudes with respect to the speed of the posterior-ward movement of the DoT, and the shear viscosity of the endoderm. To examine how these two central properties interact, we performed parametric sweeps on 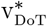 and *η*^*^ to examine the effects on tissue flows. *Q*_AP_ was highest for low 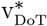 irrespective of *η*^*^, indicating that the largest tissue flows are achieved when the FGF gradient is nearly stationary (Fig 8A). As expected, increasing shear viscosity of the endoderm reduced bulk tissue flow. However, the importance of shear viscosity in this regard was largely dependent on 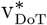, with *Q*_AP_ becoming increasingly insensitive to changes in *η*^*^ as 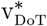 increased (Fig 8A). Notably, the experimentally measured rate of axis elongation 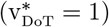 corresponds to a domain within which moderate changes in either 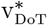 or *η*^*^ have relatively little influence on the total bulk flow, despite having more pronounced effects on maximum antero-posterior speed (Fig 8B). The anisotropy of endoderm movements was strongly dependent on both 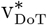 and *η*^*^, with each reducing anisotropy (Fig 8C). At sufficiently high rates of axis elongation, reducing shear viscosity produced deformations in which medio-lateral flow was larger than posterior movements (*Q*_AP_*/Q*_ML_ *<* 1) (Fig 8C).

**Figure 8.**
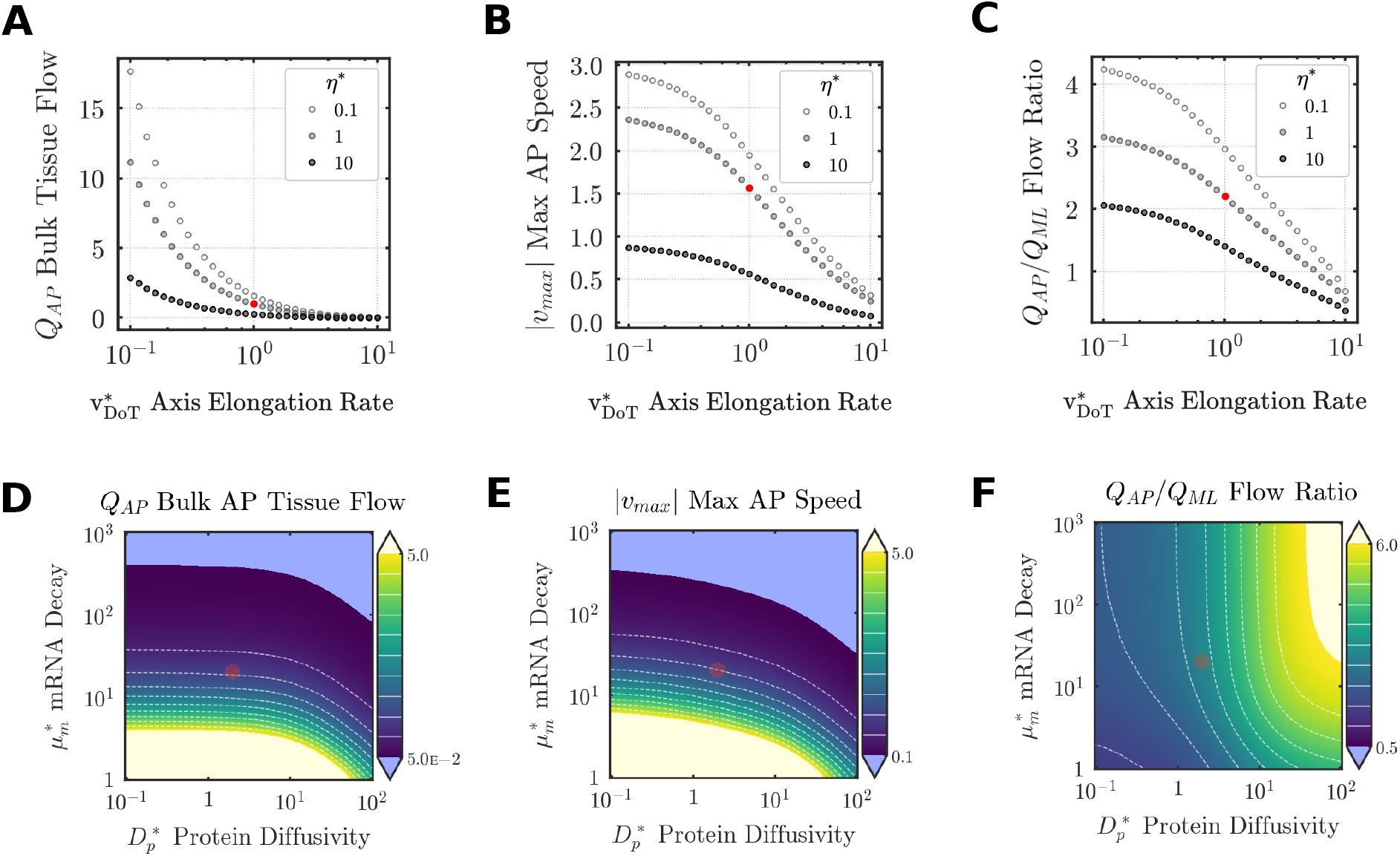
Effects of FGF transport parameters on endoderm deformations. (**A-C**) Effects of 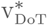 on output metrics of tissue deformation for three values of viscosity *η*^*^. (**A’-C’**) Effects of variation in 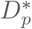 and 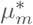 on output metrics of tissue deformation. Red circle indicates physiologic/baseline parameters as measured experimentally (**A-C**) or derived from model fits to experimental data (**A’-C’**). White dashed lines are isolines at 10% intervals of the colorbar range.

Given the importance of mRNA decay rate 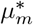 and protein diffusivity 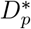 for establishment of the FGF protein gradient, we next examined how variations in each influence endoderm tissue flows. Low 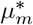 and 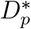, which produce an FGF gradient strongly biased along the antero-posterior axis (low *T*), also generated the largest antero-posterior flows (Fig 8D). However, bulk tissue flow was more sensitive to increases in 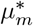 (which coordinately reduce the antero-posterior length over which the FGF gradient acts) than 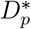, such that increases in 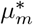 cause a pronounced decline in bulk flow as well as maximum antero-posterior velocity (Fig 8D, E). However, at sufficiently high 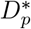, bulk tissue flow and local maximum velocity are progressively attenuated even at low 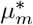. Finally, the effects of 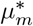 and 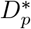 on flow anisotropy revealed a counterintuitive result: in the case of a moving FGF gradient, conditions that favor isotropic gradients (*T* = *λ* = 1) support high anisotropy, albeit for relatively small magnitudes of tissue flow (Fig 8D-F). This was entirely a result of the movement of the FGF gradient, as reducing 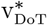 below physiologic values produces isotropic flow for isotropic FGF gradients. Together, these findings demonstrate how the mechanism of establishing a biochemical gradient is directly tied to downstream mechanics driving tissue deformations during embryogenesis.

## Discussion

In this work, we derived a chemo-mechanical model to analyze the interactions between biochemical and physical properties that drive the formation of the amniote hindgut. To do so, we began by formulating a 2-D reaction-diffusion-advection model that describes formation of an FGF protein gradient due to posterior movement of a small population of cells actively transcribing unstable *Fgf8* mRNA coupled with translation, diffusion, and degradation of resulting FGF protein. Solving the resulting pair of differential equations yields a distribution of the FGF protein concentration *p*^*^, which was used together with experimental measurements of FGF activity in the endoderm to quantify the ‘physiologic’ values of key parameters describing FGF transport. Finally, these informed a continuum model of endoderm cell movements, wherein the definitive endoderm was approximated as a viscous fluid capable of generating active tensile stresses via acto-myosin contraction in proportion to the local concentration of FGF. Parametric analyses of the resulting chemo-mechanical model provided key insights into how morphogenic and physical aspects of gut morphogenesis are integrated in the developing chick embryo.

The chemo-mechanical model replicated many key aspects of FGF gradient formation and endoderm cell movements observed during morphogenesis of the avian hindgut tube [Nerurkar et al., 2019], providing important qualitative validation for the approach. Solution of the FGF transport equations suggested that over time, the shape of the protein gradient stabilizes, such that the location of the gradient - but not its shape - changes as the embryonic axis elongates (Fig 2B). This was confirmed experimentally (Fig 3B, C), where between HH stage 10 and 15 no difference in the shape of the FGF gradient was observed despite elongation of the embryo by approximately 50% [Bellairs and Osmond, 2014]. Applying the model to experimental measures of FGF activity in the chick endoderm resulted in an approximation of FGF ligand diffusivity of 8 ∼*μ*m^2^/s. This agrees well with direct measurements of single FGF8 molecules performed in gastrulating zebrafish embryos, where a subpopulation of molecules interacting with extracellular heparan sulfate proteoglycans were found to diffuse with coefficients of ∼4 *μ*m^2^/s (compared to the faster diffusion of ∼50 *μ*m^2^/s measured in the remainder of the population) [Yu et al., 2009]. Similar diffusion coefficients have been measured for morphogens in various contexts including Dpp in the *Drosophila* wing disc [Zhou et al., 2012] and Wnt3a in the *Xenopus* gastrula [Takada et al., 2018] as well.

In conversion of this FGF gradient into endoderm deformations, the model also replicated several important experimental findings. First, as in the developing embryo [Nerurkar et al., 2019], the model revealed that FGF-induced contractility enables cell movements that outpace the rate of axis elongation (indicated by *v*_*max*_ *>* 1, Fig 7), despite axis elongation being driven by the same biochemical gradient [Bénazéraf et al., 2010, Xiong et al., 2020]. The forces associated with endoderm deformations in the model also reflected experimental observations, with the emergence of a posterior-to-anterior gradient in total stress and higher stresses observed along the medio-lateral axis than along the antero-posterior axis (Fig 6C) [Nerurkar et al., 2019]. Finally, in previous experimental investigations of endoderm cell movements during hindgut formation, we proposed that directional cell movements could be achieved from isotropic (direction-independent) cell contraction when the magnitude of contraction was spatially graded. While this was consistent with experimental observations, it could not previously be directly tested. However, in the present study, we observed computationally that, indeed, large oriented tissue flows can be generated through isotropic contraction across a range of parameter values when all mechanical properties are treated as homogeneous and isotropic.

With replication of experimentally observed behaviors spanning both FGF transport and endoderm cell movements, we next performed parametric analyses to investigate the role of various transport and mechanical properties in driving gradient formation and tissue deformations. Parametric analyses enable dissection of the system at the level of physico-chemical properties, many of which would be impractical or impossible to perturb experimentally. The relative shape and antero-posterior extent of the FGF protein gradient was most sensitive to the rate of axis elongation (Fig 4B-E), and to a somewhat lesser extent, to the decay rate of *Fgf* transcripts (Fig 5A-D). On the other hand, transport properties of FGF protein, such as the diffusion coefficient 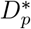, influenced the medio-lateral spread of FGF protein without a strong effect on antero-posterior gradient (Fig 5A’-D’). Given the importance of the antero-posterior FGF gradient for regulating many concurrent events in the posterior embryo, including somitogenesis [Dubrulle et al., 2001, Naiche et al., 2011], axis elongation [Bénazéraf et al., 2010, Regev et al., 2022, Xiong et al., 2020], Wolffian duct elongation [Atsuta and Takahashi, 2015, Attia et al., 2015], and hindgut tube formation [Nerurkar et al., 2019], it is possible that these findings may offer some insight into developmental constraints at play in evolution of gradient formation. Protein diffusion and stability are complex properties influenced by extracellular matrix organization and protein structure, with the latter likely constrained by the need to preserve interactions with cognate receptors and other proteins across many tissues. mRNA decay, however, can be selected upon with greater modularity as a cell-type or context-specific property. Indeed, there is growing evidence that transcript stability and post-transcriptional modification rates may explain variations in developmental timing across species [Jorge Lázaro et al., 2022], however the relative importance with respect to protein stability may be context-specific [Rayon et al., 2020]. Although morphogens have classically been assumed to form long-range gradients via diffusion from a source [Crick, 1970, Stapornwongkul and Vincent, 2021], the minimal influence of protein diffusion on establishment of the antero-posterior FGF gradient is not unprecedented. For example, the well-studied bicoid gradient that instructs antero-posterior patterning of the early *Drosophila* embryo is similarly established by a pre-pattern of bicoid mRNA (a result of active transport), with only a minor contribution from protein diffusion itself [Spirov et al., 2009]. More generally, advection as a mechanism of gradient formation has been observed in other contexts as well, whereby cell division leads to progressive dilution of mRNA as cells are displaced passively during tissue growth [Ibañes et al., 2006, Pfeiffer et al., 2000]. In these other contexts, as in the present study, it is likely that tissue movements that pre-pattern the mRNA distribution play a more prominent role in establishment of the protein gradient than protein diffusion and turnover rates.

Because the baseline transport parameters in the model were derived from experimental measurements of FGF activity in the chick endoderm, it is worth noting that the embryo occupies a neighborhood within the parameter space where the antero-posterior length of the gradient is largely insensitive to moderate changes in either mRNA or protein properties (e.g. Fig 5B, B’), but remains sensitive to the rate of axis elongation. This may explain the highly reproducible gradient shape observed from embryo to embryo, given that presumably ‘noisier’ inputs such as transcript and protein stability are less influential on the extent of the gradient than axis elongation, which is an aggregate, tissue-scale property that may be less prone to variability between individuals of a given species. It is also reasonable to expect that the rate of axis elongation will be central to gradient formation because many of the morphogenetic events that are informed by this gradient must be well-coordinated with axis elongation for proper establishment of the posterior body plan.

Extrapolation from FGF transport to downstream tissue deformation revealed several unexpected findings, and a lack of direct one-to-one correspondence between changes in transport properties and downstream readouts of endoderm movements. For example, while ellipticity of the FGF protein gradient was sensitive to both transcript decay 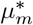 and protein diffusivity 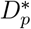 around the physiologic baseline values (e.g. Fig 5C), the resulting effects on antero-posterior tissue flow and maximum antero-posterior velocity were largely dependent only on 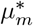. Conversely, the directionality of flow, indicated by the relative flow rates between antero-posterior and medio-lateral axes, were sensitive to variation of 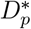 from baseline, but far less sensitive to 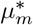. Although antero-posterior flow is the central driver of hindgut elongation, it is possible that medio-lateral movements in this population are needed during closure of the midgut tube, which is thought to rely on folding and movement of lateral cells toward the embryonic midline [Miller et al., 1999]. In this way, it is intriguing to consider that hindgut and midgut tube morphogenesis may be separable in their dependence on regulation of transcriptional and protein-level properties of the FGF gradient, respectively. However, given the paucity of studies examining the morphogenesis of midgut, it is unclear if this can be observed experimentally.

Similar to its role in establishing the gradient, the rate of axis elongation was a key determinant of tissue deformations in the endoderm as well. For example, at sufficiently high values of 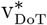, medially directed tissue flows predominated at the expense of antero-posterior movements (Fig 8C). The interaction between 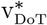 and endoderm movements is complex, owing to its influence on the system at both the mechanical and biochemical level. For example, increasing 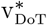T creates an elongated gradient with a lower maximum protein concentration (Fig 4B), thereby reducing the directional bias in contractility that leads to antero-posterior movement. In addition, however, a gradient that moves too quickly relative to tissue deformations will have a diminished effect on cell movements for any given gradient shape, as cells are effectively left behind as the FGF gradient moves posteriorly. These two effects together explain the precipitous drop in antero-posterior tissue movements that is observed as 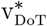 is increased (Fig 8A, B). The ability of endoderm cells to outpace axis elongation is essential for conversion of these directional cell movements into construction of a three-dimensional epithelial tube [Nerurkar et al., 2019]. Accordingly, one could interpret conditions in which *v*_*max*_ *<* 1 as unfavorable for morphogenesis and elongation of the hindgut tube. As expected, as 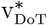 increases, ultimately cells in the endoderm are unable to keep up, suggesting that the mechanism driving hindgut morphogenesis would break down. Interestingly, however, this tipping point is dependent on tissue viscosity, a parameter that - on the long time scale of gut morphogenesis - reflects the bulk effects of dynamic cell-cell contacts, cell proliferation, and intrinsic viscosity of the cells themselves. At sufficiently high tissue viscosity, even extremely slow rates of axis elongation are not sufficient to drive hindgut formation (Fig 8B). The fastest cell movements and largest antero-posterior tissue flows are achieved in the limit as 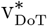 approaches zero, because this would result in the highest recruitment of cells by passive displacement of cells from low FGF concentrations to higher concentrations, where they increase acto-myosin contractility to beget even more tissue flows [Nerurkar et al., 2019]. However, operation of the system well outside of this range is consistent with the idea that this particular mechanism of gut tube morphogenesis was a secondary innovation that followed elongation of the primary body axis by FGF signaling. Indeed, mechanisms of axis elongation are well conserved among vertebrates [McMillen and Holley, 2015, Mongera et al., 2019], but the internalization of endoderm by three distinct folding events for the foregut, hindgut, and midgut, is a derived trait of amniotes alone [Durel and Nerurkar, 2020]. In other words, the posterior FGF gradient is not optimized for hindgut tube formation, but instead, hindgut morphogenesis is buffered against changes in 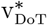, which is an external cue under distinct regulatory controls.

In the present chemo-mechanical model, FGF was only incorporated into the mechanics via a concentration dependent induction of isotropic, active tension. Therefore, this simple interaction is sufficient to fully recreate the directional anisotropic tissue deformations that drive hindgut morphogenesis, without any other sources of anisotropy or asymmetry. Nonetheless, several simplifying assumptions have been made in the present study either to improve mathematical tractability, or simply because the limited experimental studies on amniote hindgut morphogenesis leave little contextual support for more complex interactions to be considered. We have relied on downstream activity of a promoter for the FGF target gene, DUSP6, within the endoderm as a readout of FGF ligand concentration in the neighboring presomitic mesoderm, implicitly assuming that there is a proportionality between reporter activity and FGF protein concentration. The ability of the transport model to capture the 2-D distribution of DUSP6 activity when the FGF diffusion coefficient falls within the range of those reported in other contexts of FGF and other morphogens [Yu et al., 2009, Müller et al., 2012, Zhou et al., 2012, Takada et al., 2018] suggests that this may be a reasonable simplification. Further, pseudo-space analyses of published single-cell RNA Seq datasets [Ibarra-Soria et al., 2018] in the mouse endoderm suggest that FGF receptors are uniformly expressed along the antero-posterior axis (SFig 4). However, mutual antagonism between retinoic acid and FGF signaling pathways has been observed in many contexts [Aulehla and Pourquié, 2010, Lin et al., 2010], and it has not yet been studied whether the reciprocal anterior-to-posterior gradient of retinoic acid that exists in the posterior embryo [Bayha et al., 2009] has any influence on hindgut morphogenesis. From the mechanics standpoint, studies in the neighboring presomitic mesoderm suggest that FGF increases tissue ‘fluidity’ [Lawton et al., 2013], and a gradient in fluid-to-solid like physical properties has been identified along the antero-posterior axis in the presomitic mesoderm [Mongera et al., 2018]. Physical properties of the endoderm, however, have not been characterized, and heterogeneity in such properties has therefore not been observed. Cell-cell adhesion and basement membrane composition appear similar throughout the endoderm [Nerurkar et al., 2019], suggesting that any positional differences in viscosity of the cells or basement membrane may be minimal. Finally, the basement membrane was modeled as a viscous drag resisting cell movements in the endoderm, based on prior studies in the epiblast [Zamir et al., 2008], presomitic mesoderm [Bénazé raf et al., 2010], and precardiac mesoderm and foregut endoderm [Aleksandrova et al., 2015], which each suggest that the ECM in the early avian embryo is highly dynamic and tends to passively flow with cells as they reorganize. Further, disruption of contractility in the posterior endoderm during hindgut formation does not cause a full reversal of cell movements, suggesting that the basement membrane is not storing elastic energy as the endoderm flows posteriorly [Nerurkar et al., 2019]. Nonetheless, based on growing evidence of an instructive role played by the ECM during morphogenesis [Crest et al., 2017, Harrison et al., 2018], moving forward it will be important to understand how the basement membrane participates in gut tube formation. Finally, the present study only considers chemo-mechanical coupling uni-directionally, with FGF concentration influencing tissue deformations, but not vice versa. This was largely based on previous evidence that while *Fgf4* and *Fgf8* are expressed within the posterior-most endoderm, the broader expression pattern of these genes in the neighboring presomitic mesoderm aligns more closely with FGF activity in the endoderm [Nerurkar et al., 2019]. Therefore, the primary source of protein guiding tissue deformations movements is largely external to the endoderm. Nonetheless, it is possible that endoderm flows redistribute FGF proteins, and that the resulting bi-directional coupling between biochemical and mechanical properties may be important to consider as well.

The present study makes use of a relatively simple 2-D example of morphogen-mediated tissue deformations during formation of the avian hindgut to develop a mathematical model that investigates the interplay between biochemical properties that inform morphogen transport and mechanical properties that relate forces to flows in the tissue. We find that the rate of axis elongation and transcript stability are key drivers of antero-posterior movements that drive hindgut formation, while protein diffusivity and stability are limited to primarily influencing medio-lateral tissue movements. We also observed that, when constrained by experimental data defining the gradient of FGF, a corresponding graded, isotropic active tensile stress is sufficient to drive directional cell movements in the absence of any other asymmetries. A range of recent advances have expanded the toolkit available for measurement of forces, viscosity and stiffness of tissues in developing embryos [Gómez-González et al., 2020, Nelson, 2017, Davidson and Keller, 2007]. As a result, the broad goal of understanding how upstream molecular cues guide downstream mechanical effectors of morphogenesis is increasingly recognized as a central goal of developmental biology [Lenne et al., 2021]. However, given the complex interactions between biochemical and biomechanical processes, it is likely that computational approaches similar to the present study will be increasingly necessary not only for simulation, but even for interpreting experimental findings that span the molecular and physical bases of development.

## Footnotes

## Acknowledgments

We thank members of the Nerurkar Lab and Kasza Lab for their thoughtful feedback.

## Author Contributions

Conceptualization: P.O., N.L.N.; Methodology: P.O., H.C., N.L.N.; Formal analysis: P.O., H.C., N.L.N.; Investigation: P.O., H.C., N.L.N.; Resources: N.L.N.; Data curation: P.O., N.L.N.; Writing - original draft: P.O., N.L.N.; Writing - review & editing: P.O., H.C., N.L.N.; Supervision: N.L.N.; Funding acquisition: N.L.N.

## Data and Code Availability

Code for simulations can be found here (Github). Data on Dusp6 reporter are available upon request.

## Supporting Information

(Supplementary Figures 1-4, Supplementary Videos 1-2, Supplementary Methods)

**Supp. Figure 1.**
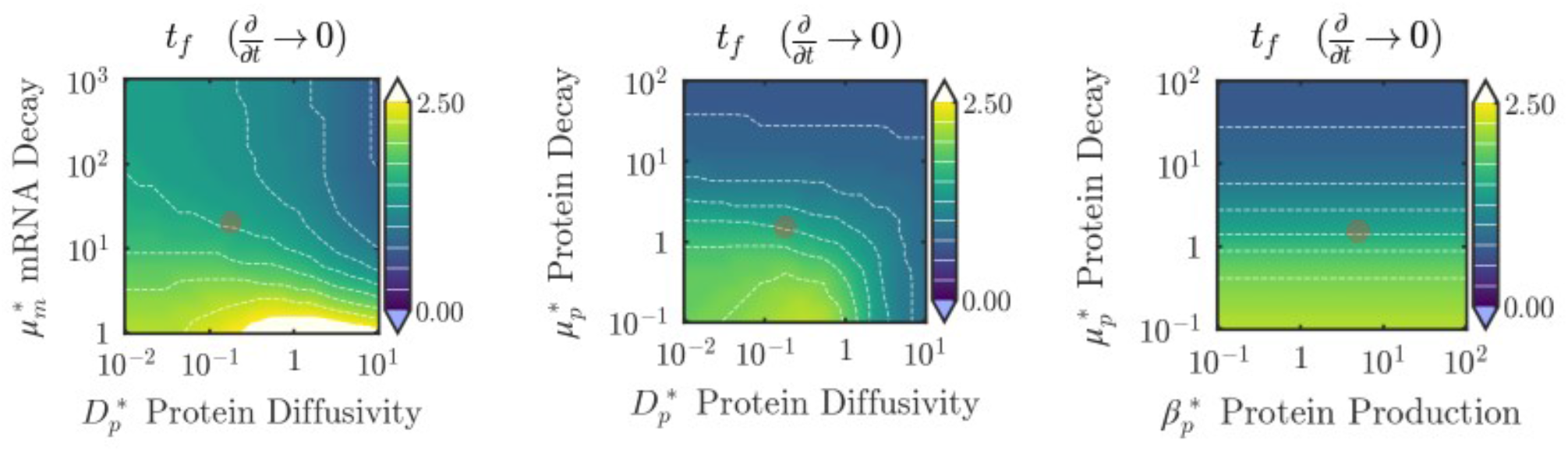
Time to steady state for the swept range of the transport parameters (all simulations on FGF transport equations were run until steady state was reached; step changes in isolines are the result of coarse time steppping, as primary focus was on steady state behavior. Red circle indicates physiologic/baseline parameter values.

**Supp. Figure 2.**
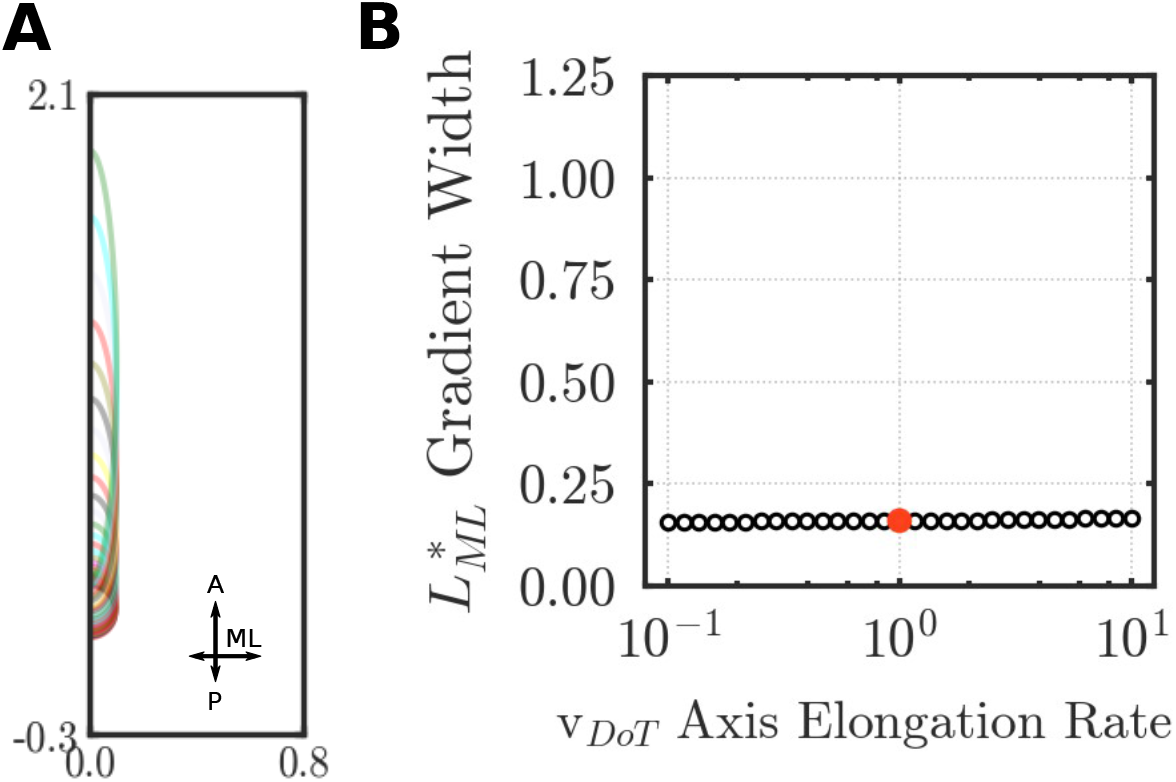
(**A**) Parametric sweep of 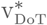 and effects on FGF gradient shape (complementary to Fig 4). Isolines indicating 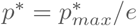 for a range of 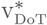 values. Isolines are pseudocolored for visibility. (**B**) Normalized medio-lateral extent 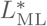 of the FGF gradient for a range of 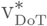 values.

**Supp. Figure 3.**
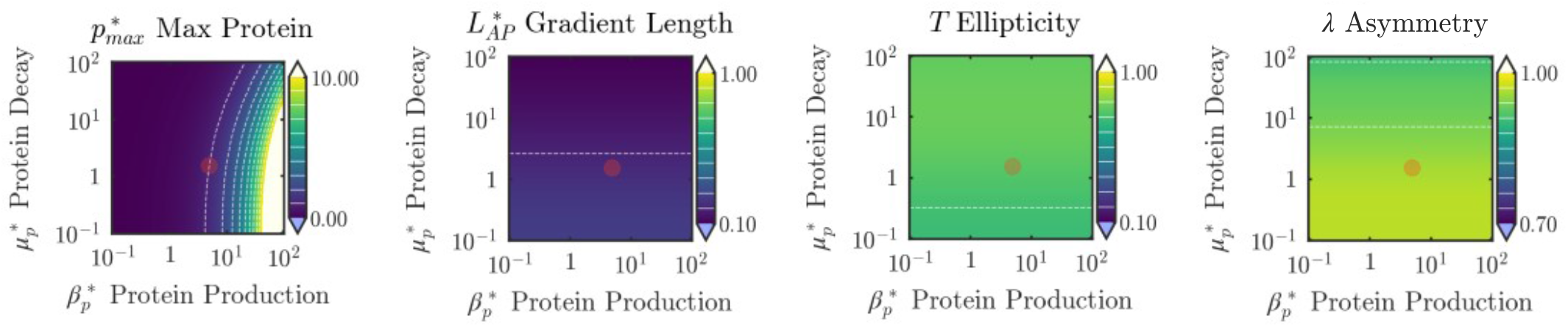
Effects of variation in protein production 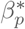 and protein decay rate 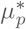 on the FGF gradient. Red circle indicates physiologic/baseline parameter values.

**Supp. Figure 4.**
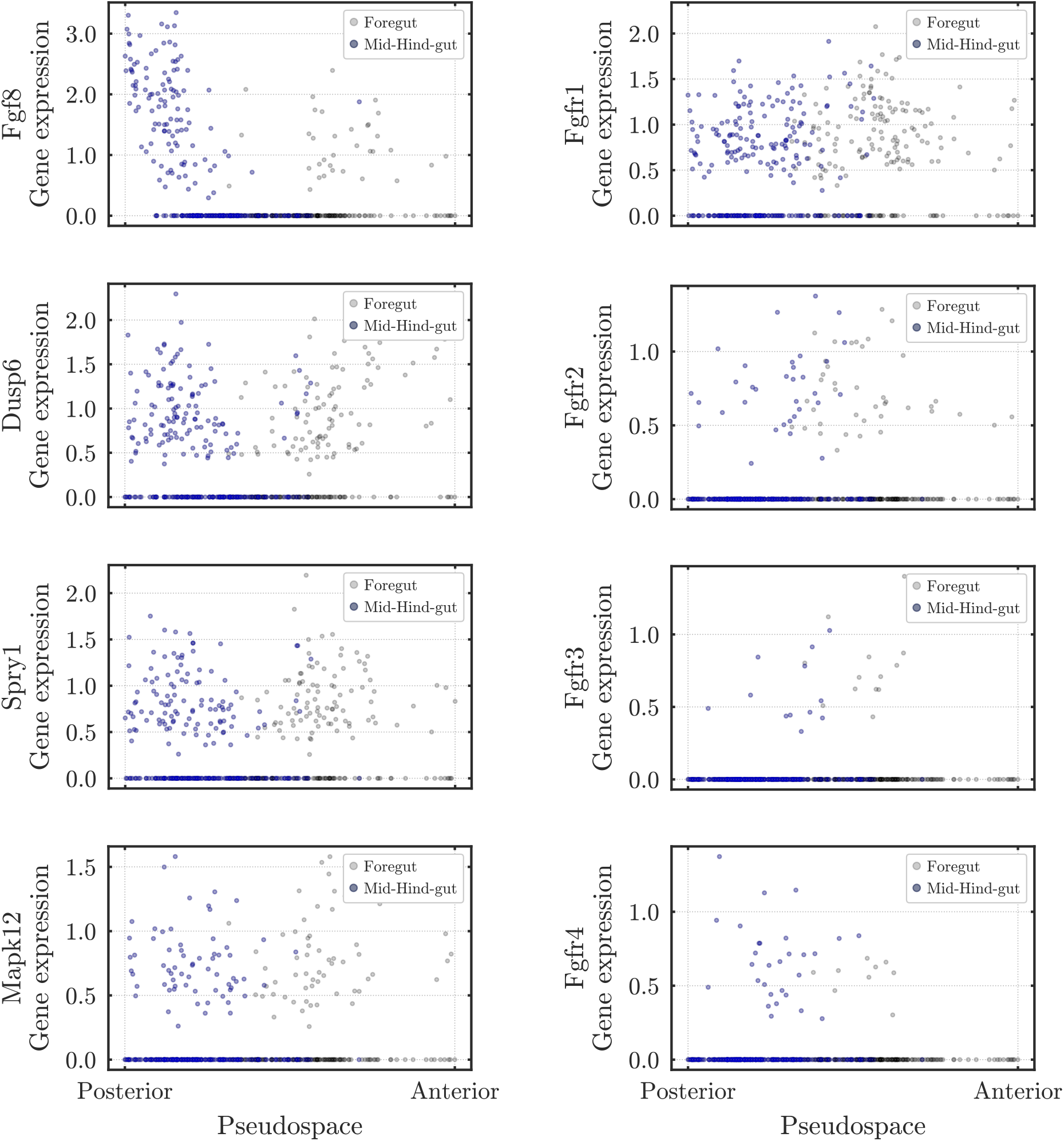
Gene expression trends (reported in log(TranscriptsPerMillion) units) per cell plotted along inferred antero-posterior axis (cells arranged in pseudo-space). Left column: FGF ligand (FGF8) and donwstream effectors (Dusp6, Spry1, Mapk12) of the pathway; right column: FGF receptors. Dataset comes from E8.25 mouse endoderm [Ibarra-Soria et al., 2018].

**S1 Video**

`mrna protein.gif`

**S2 Video**

`simulated cell tracks.gif`

## Supplementary Methods

### FGF transport - mRNA decay model

The 2-D mRNA decay model developed in the present study is of the reaction-diffusion type. In the most general form, we can write the following balance equation for the time dependent behavior of the concentration *u* for a given chemical species of interest:

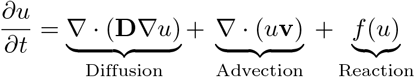

This general form of partial differential equation was used to model two species, *Fgf* mRNA (*m*) and the protein (*p*), defining the appropriate reaction term for each species. Previous studies show that *FGF8* transcription is not altered by FGF signaling, and hence there is no need for an autocatalytic term in the equations [Harrison et al., 2011].

We assume that mRNA does not diffuse, and that protein diffuses isotropically (i.e. **D**_**p**_ = *D*_*p*_**I** which simplifies the diffusion term to: *∇* (*D*_*p*_ *∇p*) = *D*_*p*_*∇* ^2^*p*. Because the model is derived and solved in a frame of reference that moves with the domain of active transcription, an advection term is introduced to account for mass flux relative to the frame of reference. The velocity of the tailbud is approximated as constant and onedimensional [Xiong et al., 2020, Bénazé raf et al., 2010], namely **v** = [*v*_*x*_, *v*_*y*_] = [0, v_DoT_], and so the advection terms for both species simplifies to 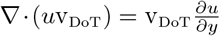 Lastly, the mRNA has a constant production rate *r*_*m*_, which is modelled as a constant 2-D Gaussian, whereas the production of the protein depends on the mRNA levels as *β*_*p*_*m*, and both mRNA and protein decay in proportion to their respective concentrations, with decay rates of *μ*_*m*_, *μ*_*p*_. Combining these terms into the generalized diffusion-advection-reaction equations above results in the following pair of coupled partial differential equations:

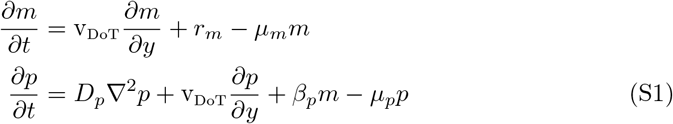

Dimensional analysis:

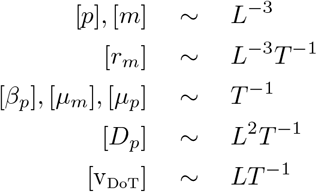

We define scaled variables:

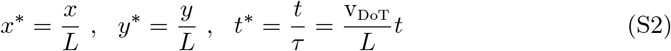

where *L* and *τ* are the characteristic length and time scale, respectively.

Equation S1 can then be written in non-dimensional form as follows

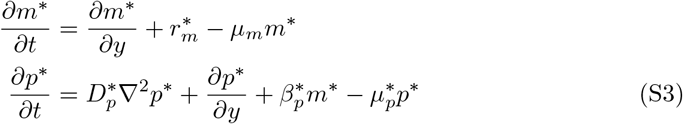

where ^*\**^ denotes a dimensionless variable or parameter. The de-dimensionalization procedure yielded the following non-dimensional parameters:

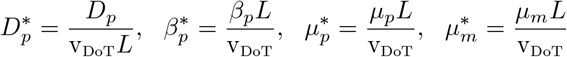

The associated boundary and initial conditions for the equations (S1) are:

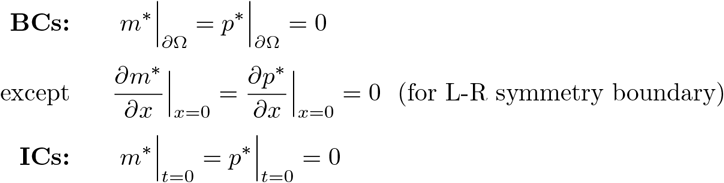

where domain Ω = [0, 2*L*] *×* [*−*2*L*, 10*L*] (antero-posterior *×* medio-lateral) and *∂*Ω is the domain boundary. N.B. we also define a subdomain of Ω, Ω_*em*_ = [0, 0.25*L*] × [0, *L*], which corresponds to the area occupied by the embryo proper (as opposed to extraembryonic tissue, e.g. hypoblast), and is where the steady-state criterion for the solution is evaluated.

v_DoT_ has been measured experimentally in the developing chick embryo [Bénazé raf et al., 2010, Bénazé raf et al., 2017, Xiong et al., 2020] ; based on these studies we used a value of 50 *μ*s/hr.

### Baseline/physiologic parameter values

The non-dimensional parameters above were estimated by fitting the model to experimental data. More specifically, to fit the protein kinetic parameters (*D*_*p*_, *μ*_*p*_, *β*_*p*_) to our experimental data (using the `scipy. optimize. curve _fit` function), the analytical solution of the 1-D reaction-diffusion-advection problem from [Harrison et al., 2011] was used:

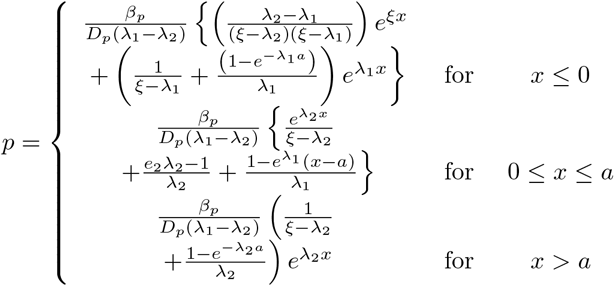

where 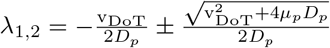, *a* = size of DoT, and 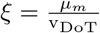

In the above equation, *p* corresponds to the protein concentration, and v_DoT_, *μ*_*m*_, *D*_*p*_, *β*_*p*_, *μ*_*p*_ are defined as above (FGF transport - mRNA decay model); for more details on solution of the 1-D problem, please consult the original publication [Harrison et al., 2011].

To extract shape descriptors of the FGF gradient, an isoline at 1*/e* of the maximal concentration was fit to the ovoid function [Baker, 2002], using the `scipy.optimize.curve_fit` function. The ovoid function offers a simple formula to approximate asymmetric elliptical shapes (similar to those observed experimentally), with the ellipticity *T* encoding the shape aspect ratio, and the asymmetry *λ* the reciprocal blunting of one end and sharpening of the opposing end of the shape.

Before the 2-D FGF-transport equations can be solved using the Finite Element Method, they must be cast in their weak form, derived by multiplying by a test function *ψ*, integrating over the entire domain, and resolving any second or higher order derivatives using the product rule of integration and ‘off-loading’ the derivatives to the test function [A. Logg et al., 2012]. The only term with a second order derivative (i.e. which requires special handling) is in the equation for the protein concentration, which was treated as follows:

1. Multiply by test function *ψ*

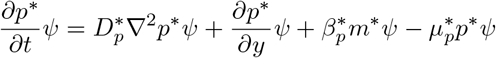
2. Integrate over Ω

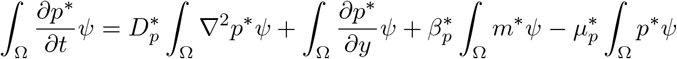
3. Use Integration by parts - this part only pertains to the *diffusion* term, where the Laplacian operator (*∇*2) appears,

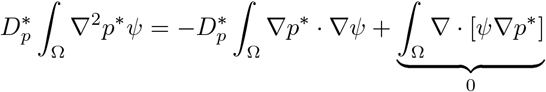 Conversion of the final integral above into a surface integral by way of Green’s Divergence Theorem results in a term, 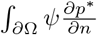, which vanishes owing to the no flux boundary conditions, i.e.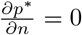 on *∂*Ω.
4. Discretize time derivative operator using first order backward Euler scheme:

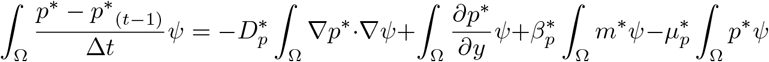

where *p*^*\**^(*t−*1) is the solution from the previous time step (starting with the initial condition).

The model was solved using the python-based `FEniCS` finite element solver by directly inputting this weak form.

### Tissue Mechanics - Active fluid model

To model endoderm mechanics, we use the theory of active fluids - the epithelium is modeled as a viscous fluid, with active tensile stress modulated by the concentration of FGF protein obtained from solving the FGF transport model described above.

On the long time scales over which morphogenesis occurs, inertial effects can be ignored so that local force balance reads

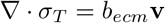

where the total stress *σ*_*T*_ = *σ*_*P*_ + *σ*_*A*_ is the sum of the passive stress and the active stress.

The active stress is fully isotropic and reads:

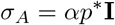

where *p*^*\**^ is the FGF concentration as computed by the FGF transport model above and *α* is a factor to convert from units of concentration to units of stress.

The passive stress can be further decomposed as:

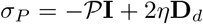

where 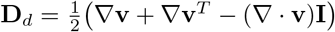**; v** = [*v*_*x*_, *v*_*y*_] is the velocity, is the pressure and *η* the viscosity.

We model the fluid as compressible, to permit area changes in 2-D that are accommodated by out of plane movements (i.e. allow negative divergence) [Serra et al., 2021]:

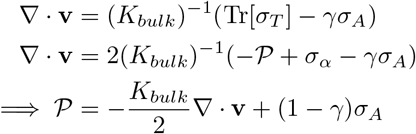

where *K*_*bulk*_ the bulk viscosity and *γ* a factor that controls the amount of out of plane motion accounted for by the active stress.

Lastly, we model the interactions with the ECM on the basal surface as a viscous drag force *b*_*ecm*_**v** where *b*_*ecm*_ is the viscous drag coefficient.

Putting this all together, the force balance equation reads:

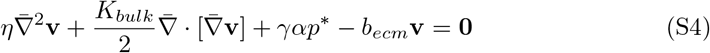

Dimensional analysis:

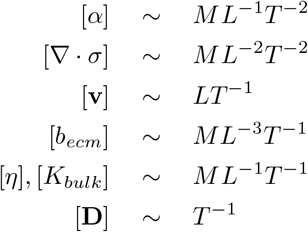

These equations S4 can be written in non-dimensional form as follows

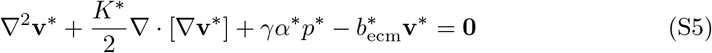

where ^*\**^ denotes a dimensionless variable or parameter; Using the same scaling as in S2, we define the following non-dimensional parameters:

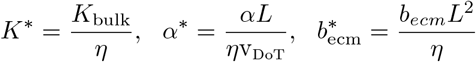

The associated boundary and initial conditions for the equations (S5) are:

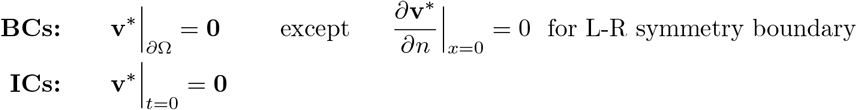

where domain Ω = [0, 2*L*] *×* [*−*2*L*, 10*L*] and *∂*Ω is the domain boundary.

Similar to before, we need to cast equations S5 in their weak form, following a similar procedure.

### Single-cell RNA-sequencing data analysis

A previously published and publicly available data set [Ibarra-Soria et al., 2018] comprising stage E8.25 embryonic mouse endoderm was analyzed to examine the spatial distribution of FGF related genes along the antero-posterior axis. Data was accessed using the online tool (marionilab.cruk.cam.ac.uk/organogenesis, Accessed 2022-10-10). Analysis was carried out with the Python package `scanpy` [Wolf et al., 2018], part of scverse. A diffusion-based method (`scanpy.tl.dpt`) was used to arrange cells in antero-posterior pseudo-space, validated by region-specific (fore-, mid-, hind-gut) expression patterns.

## Code Availability

The code to solve the equations of the chemo-mechanical model presented above is deposited in the form of an executable ‘notebook’ (Github). Please consult the official FEniCS website for installation instructions.

